# Transcriptome Analysis of Archived Tumor Tissues by Visium, GeoMx DSP, and Chromium Methods Reveals Inter- and Intra-Patient Heterogeneity

**DOI:** 10.1101/2024.11.01.621259

**Authors:** Yixing Dong, Chiara Saglietti, Quentin Bayard, Almudena Espin Perez, Sabrina Carpentier, Daria Buszta, Stephanie Tissot, Rémy Dubois, Atanas Kamburov, Senbai Kang, Carla Haignere, Rita Sarkis, Sylvie Andre, Marina Alexandre Gaveta, Silvia Lopez Lastra, Nathalie Piazzon, Rita Santos, Katharina Von Loga, Caroline Hoffmann, George Coukos, Solange Peters, Vassili Soumelis, Eric Y. Durand, Laurence De Leval, Raphael Gottardo, Krisztian Homicsko, Elo Madissoon

## Abstract

Recent advancements in probe-based, full-transcriptome, high-resolution technologies for Formalin-Fixed Paraffin-Embedded (FFPE) tissues, such as Visium CytAssist, Chromium Flex (10X Genomics), and GeoMx DSP (Nanostring), have opened new opportunities for studying decades-old archival samples in biobanks, facilitating the generation of data from extensive cohorts. However, the experimental protocols can be labor-intensive and costly; therefore, it is thus essential for researchers to carefully evaluate the strengths and limitations of each technology in relation to their specific research objectives.

Here, we report the results of a comparative analysis of the three methods mentioned above on FFPE archival tumor samples from four non-small cell lung cancer, four breast cancer and six diffuse large B-cell lymphoma. We highlight some relative advantages and disadvantages of each method in the context of operational challenges, bioinformatic analysis and biological discovery. Our results show that: 1) all three methods yielded good-quality, highly reproducible transcriptomic data from serial sections of the same FFPE block; 2) GeoMx data contained mixtures of cell types, even when pre-selecting areas with cell type-specific markers; 3) high-throughput spot-level (Visium) or cell-level (Chromium) data enabled the identification of tumor heterogeneity within and between patients, which could be used to identify targeted therapies.

Our data support the use of Visium and Chromium for high-throughput and discovery-driven projects, while the GeoMx platform could be suited for addressing specialized questions on targeted regions. All data generated from this study, including GeoMx, Visium, Chromium, H&E, and expert annotations are publicly available.

## Introduction

Recent advances in sequencing- and imaging-based techniques have led to the development of spatially resolved transcriptomics, named method of the year in 2020 ^1^ with the ability to spatially quantify gene expression within a given tissue. These technologies allow health researchers and clinicians to characterize patient samples with unprecedented depth and spatial resolution, leading to transformative insights that have the potential to improve diagnoses, treatments, and patient outcomes.

Hybridization-based full-transcriptome methods relying on short probes have proven successful on FFPE tissue: Visium v2(Visium)^2^, Chromium Flex (snRNAseq)^3^ and GeoMx DSP (GeoMx)^4^ methods profile 22,000 transcripts on the human genome, including the majority of the protein-coding transcripts necessary for the discovery of novel biological mechanisms and new drug targets. Visium v2 enables uniform coverage of tissue with ∼5,000 of spots of 55 µm diameter, spaced 100 µm apart. GeoMx DSP allows users to pre-select regions of interest (ROI) on the tissue, and collect spatially distributed segments identified by fluorescence markers called areas of illumination (AOI). Chromium Flex is a single-cell/ single-nuclei transcriptomics platform with similar performance to the previous droplet-based methods, adapted for FFPE tissues^5^. Each individual method is both laboursome and costly to set up and perform. The relative strengths and weaknesses of the methods need better assessment to inform researchers on which technology to use.

There have been attempts at comparing Visium v1 (version 1) and GeoMx DSP. Technical performance in similar tissues showed higher sensitivity but lower specificity on gene detection for GeoMx, using comparative experimental design for both methods^6^. Since then, Visium was updated with an automated sample transfer tool, Cytassist, which is considered to yield even higher sensitivity and specificity compared to Visium without Cytassist (version 1), making this a method of choice for the current study^7^. Following the release of single-cell RNA-seq in FFPE tissue on Chromium, the combined use of Visium and Chromium effectively demonstrated an improved performance for identifying and mapping malignant cell subtypes within a single patient^3^. All three methods were used together to build an atlas of cholestatic liver disease: Chromium for determining cell states, Visium for determining tissue regions, and both Visium and GeoMx for confirming co-localisation of cell types in regions^8^. While all the methods have been applied in cancer samples, there is a lack of consensus in the field of which method is best suited for high-throughput discovery projects for hundreds of patients. To bridge this gap and to guide technology selection, we performed a comparative study between three full-transcriptome methods with GeoMx, Visium and Chromium.

Here, we aimed to study the operational efficacy, identify complementarities, and propose optimal use-cases for each of these three technologies while taking advantage of the strength of each method. We analyzed 14 patients with all three methods from FFPE blocks across three cancer types: breast cancer, non-small cell lung cancer, and Diffuse Large B-Cell Lymphoma (DLBCL). We assessed the cell type signature specificity between the spatial transcriptomics (ST) methods, explored tissue heterogeneity and demonstrated intra- and inter-patient variations that can inform precision therapy decisions. Our study highlighted the strengths, weaknesses, and complementarity of each approach, and recommended Visium and Chromium for high-throughput discovery projects.

## Results

### Experimental overview and data curation

First we generated a dataset of spatial and single-nuclei transcriptomes across three cancer types. Consecutive tissue sections of archival FFPE blocks (median age 57 months (22-103); median DV200 53.75% (7.2-80.3)) of breast) and lung cancer resections (n=4 each) and diffuse large B-cell lymphoma (DLBCL) excisional and sampling biopsies (n=3 for each) were submitted for profiling with GeoMx, Visium v2 and Chromium FLEX platforms (Figure 1a, Supplementary Figure 1a, Supplementary Table 1-3). Tissue slices from each specimen with maximum size of 11mmx12mm were placed on GeoMx slides by groups of three and stained with fluorescence markers to collect the corresponding AOI-s (Figure 1a, Supplementary Figure 1a). For the Chromium FLEX workflow four samples were pooled in a single well. The nuclei were prepared using the snPATHO protocol^9^ from two 25 uM cuts and were used for single-nuclei preparation. Wide distribution of different RNA quality values with DV200, histology types and block ages were chosen (Figure 1b, Supplementary Figure 1a).

**Figure 1:**
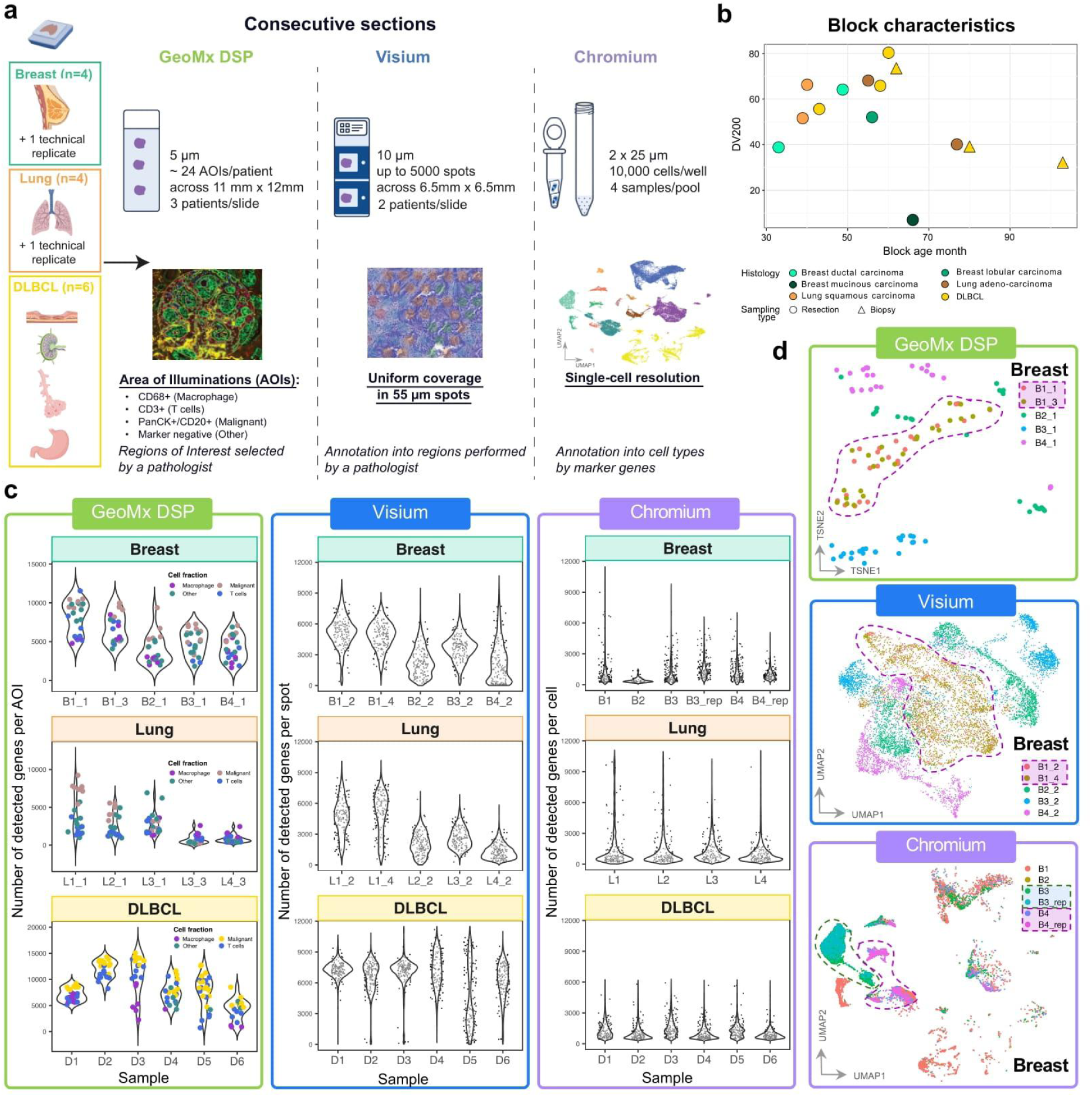
Experimental setup and data quality with three full transcriptome methods GeoMx, Visium and Chromium on Breast and Lung cancer, and DLBCL biobank samples. (a) Schematics of experimental design to generate GeoMx and Visium spatial transcriptomics and Chromium single-nuclei data for three cancer types corresponding to 14 donors from FFPE blocks with 2 replicates for each technology. AOI - area of illumination, DLBCL - Diffuse large B-cell lymphoma, DSP - digital spatial profiler (b) Display of block age and RNA quality measure DV200 values for all the samples with histology type. (c) Gene detection rate for GeoMx AOI-s and number of genes detected in Visium spots and single nuclei. Colors in GeoMx correspond to AOI labels. (d) Breast cancer tSNE plots with GeoMx and UMAP plots with Visium and Chromium show good reproducibility. Technical replicates with adjacent sections from the same block are highlighted with a dashed line. B - breast, L - lung, D - DLBCL.

The number of data units varied across methods and samples (Supplementary Figure 1c-e). GeoMx AOI-s were successfully selected for about 24 AOI-s per patient. Number of spots in Visium varied from 822 in a Breast cancer sample to 4951 in DLBCL depending on the size of the tissue. Chromium recovery was more variable from 802 nuclei to 17,804 nuclei per sample. We also observed high variation in the number of transcripts and in the number of genes detected across the samples with all three methods (Figure 1b, Supplementary Figure 1b).

To assess reproducibility, we compared data from adjacent sections in Breast (Figure 1d). The samples performed consistently both in gene detection, as well as by clustering using high-dimensional representation. We observed a slide effect between replicates of lung samples in GeoMx, which was later corrected in downstream analysis (Methods). Overall, we conclude that the sample-level variability in data dominates the variability induced by processing, demonstrating high reproducibility for all three technologies. Overall, we present a highly reproducible spatial and single nuclei dataset characterizing the transcriptome of archival samples from three cancer types.

### GeoMx data exhibit non-specific signals in pre-selected AOI segments

We first aimed to compare the ST methods GeoMx and Visium for their capacity to characterize cell types in tissue. For this purpose, we used image-based (i.e. H&E derived) labels for Visium spots and manually annotated Chromium data. AOI segment identity was used for GeoMx (Supplementary Figure 1g, Supplementary Table 4a-c). For Visium, pathologists used the Loupe Browser (10X Genomics) to annotate all spots into tissue categories based on H&E images. These categories were then grouped into broader classifications: Tumor-enriched, Stroma-enriched, Lymphocyte-enriched, Immune Cell Mix (lung and breast), and Epithelia (DLBCL gastric and bronchial biopsies) (Supplementary Figure 1f, Supplementary Table 5). The single-nuclei data was annotated into known cell types according to previously described markers and assigned to four hierarchical levels of cell type groups (Figure 2a, Supplementary Figures 2, 3, Supplementary Table 6-7). Single-cell annotation at highest resolution (Level 4) enabled the identification of subtypes of malignant cells within one donor, effectively capturing intra-patient heterogeneity.

**Figure 2:**
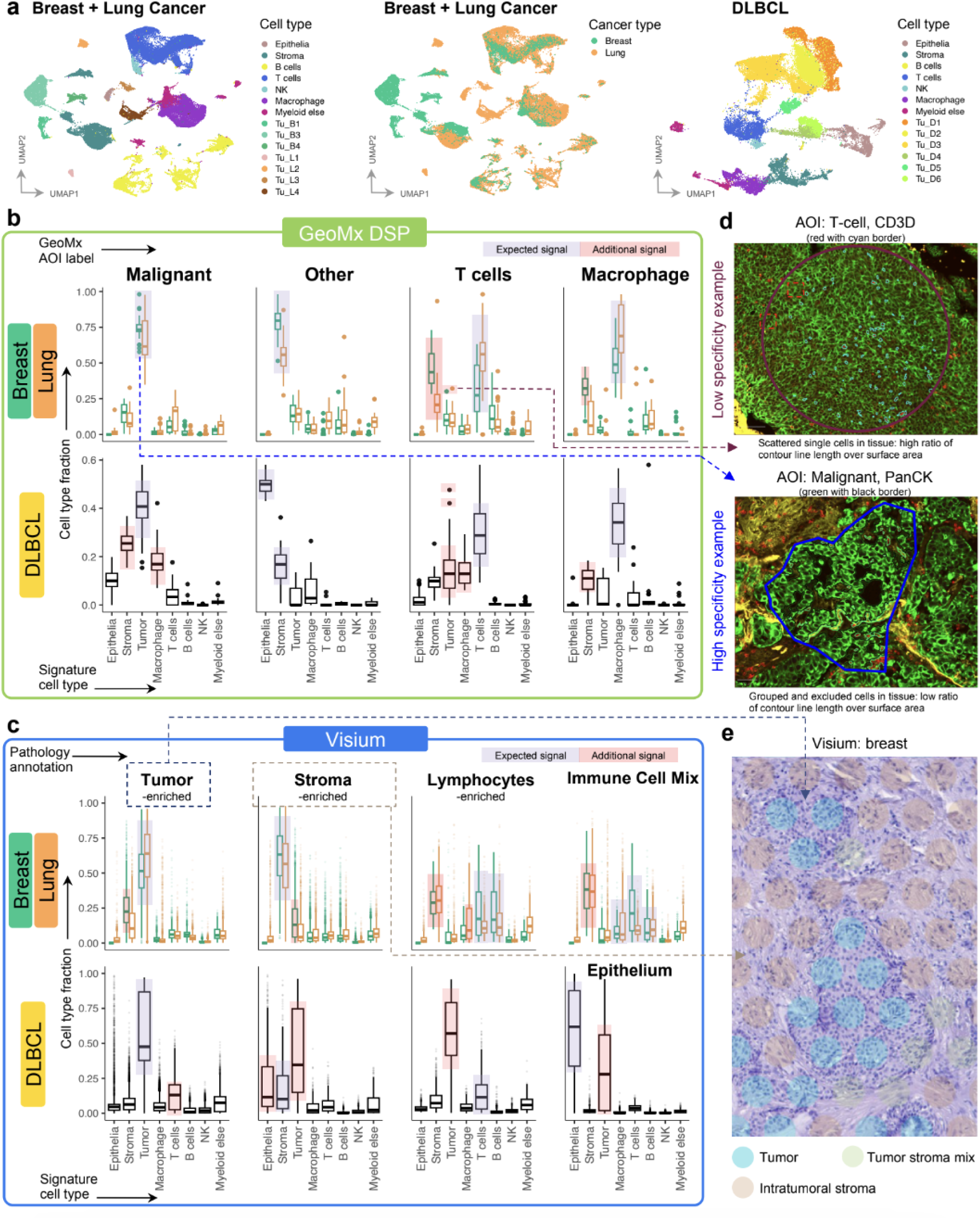
Assessment of cell type specificity in GeoMx AOI-s and Visium spots. (a) UMAP plots for annotated Chromium data at Level 1.5 resolution. Predicted cell type fractions by deconvolution in GeoMx (b) and Visium (c), grouped by AOI label in GeoMx and Pathologist annotation groups in Visium. Fractions are displayed for groups of shown cell types. Expected and unexpected additional signals for that AOI label or pathology group is highlighted. Pathology annotations were grouped as in Supplementary Table 5. (d) Examples of AOI-s with low and high specificity on immunofluorescent images. Thicker line is contour for the full region of interest, thinner line is contours for AOI. (e) Examples of Visium spot annotation into Tumor and adjacent areas.

The cell-type composition of the AOI-s and spots was then derived by deconvolution of each data point using our Chromium reference data with the SpatialDecon^10^ tool for GeoMx and Cell2location^11^ for Visium (Methods). Both Visium and GeoMx methods captured enrichment of expected cell types in agreement to the Visium pathology annotation and the GeoMx curated AOI label (Figure 2b, c). Also, in both methods tumor and stromal cell types were predominantly enriched in their respective Tumor and Stromal regions. T-cell signatures were enriched in *T cells* AOI-s in GeoMx, and the T- and B-cell signatures were enriched in Lymphocyte regions in Visium. *Macrophage* AOI-s in GeoMx and Immune cell mix in Visium also matched expected cell type enrichments. In addition to the expected signal, there was an unexpected stromal signal for all the immune-related regions in both methods. This was expected in Visium where, by design, spots capture around 20 cells. In GeoMx, nonspecific signal in AOI capture was also observed with canonical markers’ expression in Breast, Lung, and DLBCL (Supplementary Figure 4a, b). The cell type specificity on a global scale was lost in the Breast *T cells* that clustered together with *Other* segments (Supplementary Figure 4c). Although the GeoMx method was designed to capture single cell types, we observed that the least specific signals originated from AOIs containing scattered cells, such as *T cells*. In contrast, the purest signals were obtained from tissue areas with high uniformity and a large surface area relative to the AOI contour (Fig 2d). The nonspecific signal from Visium is explained by the spots laying at intersection between two areas, and cell type mixtures positioned within the same area (Fig 2e). Additionally, our H&E-derived labels assigned spots based on their predominant cell type(s), so background signals from other cell types were expected (Methods). Overall, we concluded that both spatial methods captured mixtures of cells that can be readily deconvolved computationally. However, in the case of GeoMx, the selection of AOIs within cell clusters versus scattered cells in the tissue can influence the specificity of the signal.

### Cell type specificity in matched GeoMx and Visium regions

To directly compare GeoMx and Visium, we aligned images to match Visium spots with specific AOI locations in GeoMx (Methods, Figure 3a). In total, nine pairs of consecutive sections of GeoMx and Visium breast and lung samples were used for alignment (Supplementary Figure 5a, d). When 70% of the Visium spot’s area was overlapping with a GeoMx segment, it was considered matching to that segment’s AOI label. Around 11% of the spots that fell into an AOI got spatially mapped to an AOI label. *Malignant* and *Other* AOI-s had the highest number of spots mapping to their area providing 116 and 342 data points respectively (Figure 3b). Spots mapping to either *T cells* or *Macrophage* AOI-s had a smaller overlapping area to a GeoMx segment, providing only five spots for *T cells* and no matching spot for *Macrophage* AOI label. This is due to the gaps between Visium spots, and the irregular shapes of immune cell marker staining in GeoMx that fail to be captured by Visium spots. For the location-matched spots, we compared the cell type specific signal with deconvolution between GeoMx and Visium (Figure 3c). Both technologies showed comparable cell type fraction in *Malignant* and *Other* AOI-s. In matched *T cells* AOI-s, we saw a higher proportion of stroma cells than T cells in GeoMx, while such a background signal is suppressed in Visium. Note that all 5 mapped spots belonged to breast samples (Supplementary Figure 5d), for which the stromal signal is lower than T-cell signal compared to in lung samples (Figure 2b). To estimate the signal from these minority cell types in GeoMx and Visium independently of location matching, we display the T-cell and macrophage deconvolution fractions of each AOI or spot on integrated reduced dimension plots for all indications (Supplementary Figure 7a, b). Lung samples show higher abundance of T-cell and macrophage spots in Visium, consistent with cell type proportions in Chromium (Supplementary Figure 3). However, we observe a higher deconvolution fraction of T-cell and macrophage in GeoMx’ manually selected AOI-s (Supplementary Figure 7b). This demonstrates the strength of highly segmented GeoMx design that allows a better enrichment for rare cell types of interest in a tissue.

**Figure 3:**
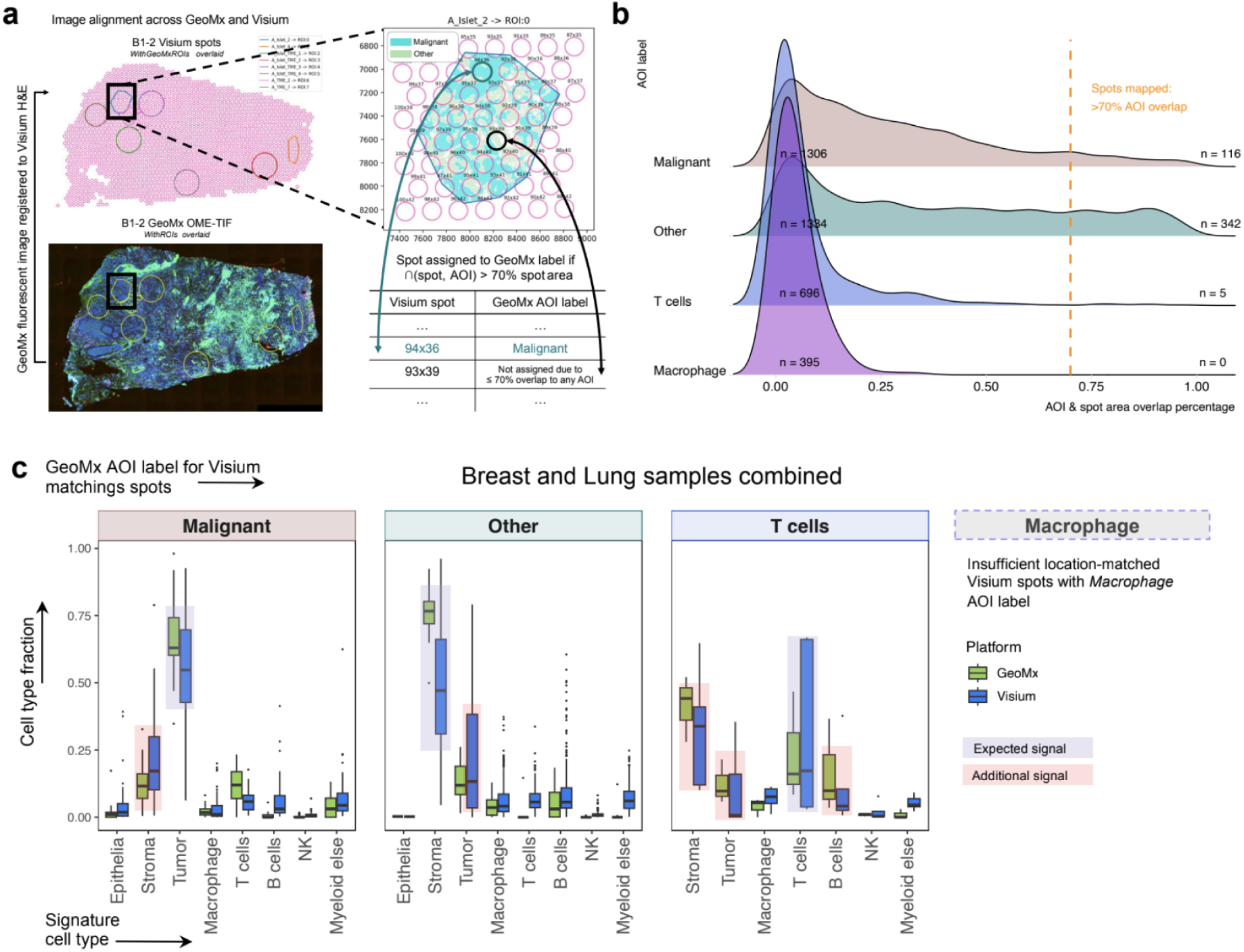
Cell-type deconvolution specificity comparison on spatially registered Visium spots to GeoMx AOI labels. (a) Image registration between GeoMx fluorescent image and Visium spot coordinates. (b) Density plot of overlapping area between Visium spots and each GeoMx AOI. A threshold of >70% was applied to select spots registered to a GeoMx AOI label. (c) For registered GeoMx segments and Visium spots in (b), using matched Level 4 Chromium reference from Fig 2a to deconvolute cell type fraction.

Overall, we conclude that Visium and GeoMx demonstrate comparable specificity in head-to-head spatial registration comparisons and effectively capture signals from rare cell types. Notably, despite the enhanced cell-type enrichment offered by GeoMx’s AOIs, Visium reliably detects signals from corresponding regions.

### Spatial profiling adds resolution to the H&E-based annotations in Visium

Next, we explored data-driven spot-level annotations instead of relying on pathologists’ H&E-based annotation in Visium. The deconvolution of each spot into cell type fractions enables higher resolution for visualization of the tissue into cell-types that are impossible to distinguish based on H&E alone, as demonstrated for sample B1 (Figure 4a, b). Deconvolution also agreed with the pathologist’s observations, as evidenced by the breast, lung (Supplementary Figure 5b, c) and DLBCL (Supplementary Figure 6) patients data. The increased resolution is evident in regions with similar histology, but different cell type composition. First, T- and B-cell were annotated as Lymphocytes on H&E, but deconvolution distinguished these cell type groups as expected (Fig 4c). Second, malignant cell subtypes identified in Chromium for patient D3 showed distinct localisation on tissue (Fig 4d). Finally, neoplastic B-cell and stomach epithelium layers in DLBCL biopsy are clearly separated by cell type signatures into distinct regions, although intermixed with pathology annotations (Fig 4e). Overall we demonstrate that data-driven annotation provides increased resolution compared to the H&E-based manual annotation.

**Figure 4:**
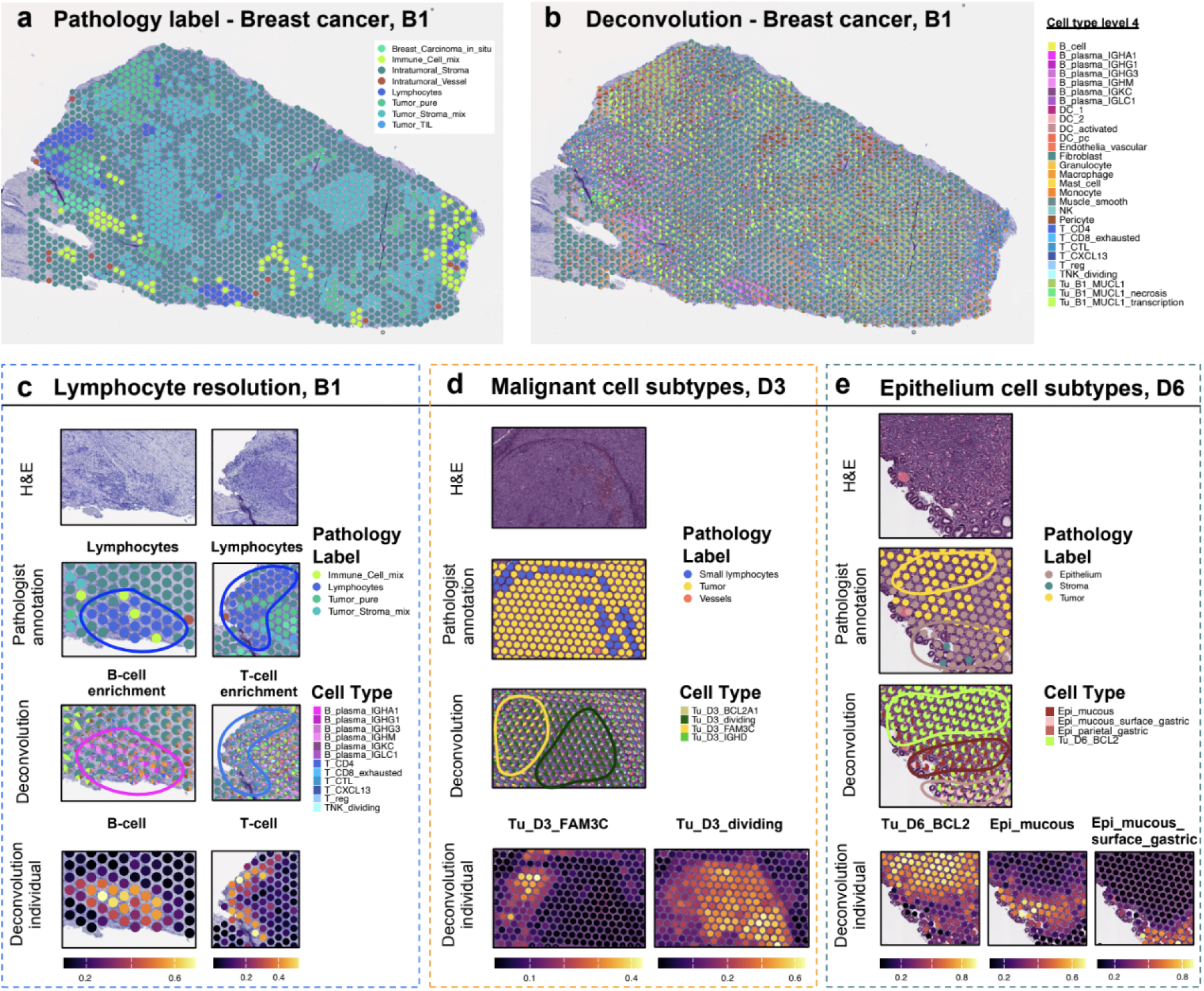
Multi-modality of Visium data visualized with H&E image, pathology label, and deconvolution. (a) Pathology label visualized on H&E image of sample B1_4. (b) Deconvolution cell type fraction plotted as scatterpie for each spot visualized on H&E image of sample B1_4. (c) Lymphocyte resolution zoomed in on H&E image, pathology annotation, deconvolution shown as scatterpie of cell type fractions, and individual deconvolution fraction for cell type of interest in sample B1_4. (d) Malignant cell subtypes visualized across various modalities for sample D3. Deconvolution is able to depict the spatial transitioning of tumor cell types annotated in Chromium in Visium. (e) Epithelium cell subtypes visualized across various modalities for sample D6.

### Spot-level data can be enhanced computationally to increase resolution and detection rates

A common criticism of the Visium technology is its resolution and data sparsity^1^. Each spot typically contains an average of about 20 cells, and similar to single-cell/nuclei RNA-seq, the expression levels for canonical marker genes can be relatively low, making it challenging to identify and visualize detailed tissue structures. To address this limitation, we applied BayesSpace^12^, a reference-free approach that subdivides each spot into subspots, estimating the gene expression contribution of each subspot to the overall spot-level value, thereby generating a super-resolution image (Figure 5). With its high-resolution gene expression maps, BayesSpace can resolve tissue structures that are difficult to detect at the original resolution. To illustrate this, we analyzed a sample containing a visible tertiary lymphoid structure (TLS) on the H&E image. The corresponding area in Visium shows limited evidence of TLS, with sparse expression of canonical marker genes such as CXCL13, MS4A1, and CD4 (Fig 5a). However, after enhancement with BayesSpace, the same region reveals a well-defined area with elevated expression of these marker genes. These findings show that the resolution of Visium data can be significantly enhanced, thereby increasing its analytical value for spatial profiling.

**Figure 5:**
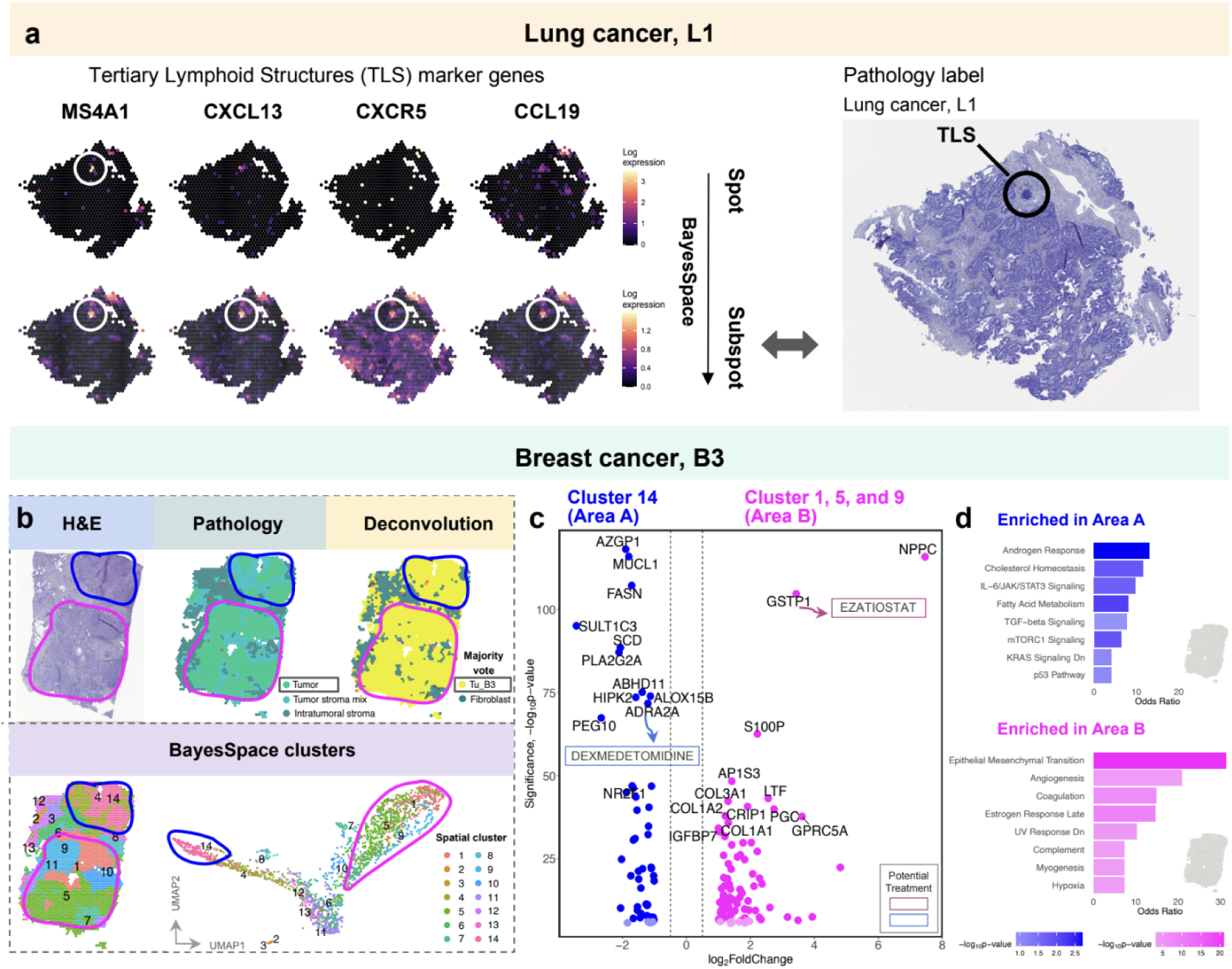
Integrating H&E image, pathology label, deconvolution, and clustering identifies TLS and tumor subtypes in Visium. (a) TLS markers show clear boundary of TLS after BayesSpace resolution enhancement, which validates the TLS location by pathology label. (b) Two regions similar in H&E, labeled as tumor by pathologist, deconvolution majority voted as single tumor type from Chromium: two spatially distinct tumor subtypes appeared with spatially-aware clustering. (c) Volcano plot of Differential Expression (DE) analysis between selected clusters from b. Known drug targets and their corresponding drugs are highlighted. (d) Gene set enrichment analysis for the DE genes between the two tumor areas from c.

### Visium identifies spatially disjoint malignant regions with different transcriptional profiles

In addition to identifying structures, we aimed to look for intra-patient variation in Visium. From the H&E image of B3, there was no morphological difference between the two highlighted regions, annotated as “Tumor_pure” (Fig. 5b). Cell type majority vote from deconvolution analysis confirmed that spots in these areas are mostly labeled as “Tu_B3”. Using the spatially-aware clustering method, BayesSpace, we discovered that the two areas are transcriptionally distinct on the reduced dimension UMAP. We group the clusters as cluster 14 (Area A) and clusters 1, 5, and 9 (Area B). Differential expression (DE) analysis between the two tumor areas of Area A against Area B showed distinct profiles. Some of the DE genes identified are drug targets, for example, a potential treatment for marker GSTP1 in Area A is ezatiostat and for marker ADRA2A in Area B is dexmedetomidine. Gene set enrichment analysis between the areas showed a mixture of signaling pathways, such as^13^ IL-6/JAK/STAT3 signaling^14^, mTORC1 signaling^15^, and p53 pathways that show signs of tumor proliferation, and pathways such as KRAS signaling down^13^ and TGF-beta signaling^16^ suggesting tumor suppression^16^ and immune infiltration. In Area B, we see enrichment of Epithelial mesenchymal transition (EMT) and angiogenesis pathways that contribute to more aggressive behavior of cancer cells^17^. Interestingly, Area B had additional signal from Fibroblast cell types by deconvolution, which suggests the interplay with fibroblasts in more aggressive tumor areas (Supplementary Figure 7c). This intra-patient heterogeneity was replicated with Chromium data where we annotated two tumor subclusters for Tu_B3 corresponding to the area A and B specific DE genes PLA2G2A and NPPC correspondingly (Supplementary Figure 7d). The two subclusters mapped to the expected areas in Visium, and the DE analysis showed similar significant DE genes as the spatial analysis (Supplementary Figure 7e, f). We observed an agreement of IL-6/JAK/STAT3 signaling pathways in Area A between Chromium and Visium. However, with enhanced resolution of the tumor subcluster in Chromium, Area B also displayed more detailed pathways of tumor progression (e.g. mTORC1 signaling^15^) and immune regulation (e.g. Interferon alpha response^18^) (Supplementary Figure 7g). This discovery highlights the independent potential of both Visium and Chromium to uncover intra-patient heterogeneity that standard pathology using H&E image-based annotation may miss. Notably, Visium also offers the capability to spatially map these subtypes.

### Potential for targeted therapies in heterogeneous patient population

Besides molecular characterisation of intra-patient heterogeneity in Breast, we wanted to explore the power of full transcriptome methods to describe existing drug targets and discover new ones in a highly heterogeneous and complex tissue such as DLBCL. We first aggregated annotations of DLBCL malignant and tumor microenvironment (TME) cells to similar categories (Figure 6a) by using Level 2 categories in Chromium with malignant subtypes, annotating Visium spots and using GeoMx AOI labels. After correcting for donor effect, Visium annotation into epithelium, stroma, plasma, and vessels/immune was done using clustering and known marker gene enrichment (Methods, Supplementary Figure 8). We used canonical marker expression and relied on pathologist annotations for Necrosis regions. Malignant nuclei were annotated by both enrichment of patients in the cluster as well as the cell type signature similarity with the corresponding donors’ malignant cell markers gained from Chromium data (Supplementary Figure 3b-d).

**Figure 6:**
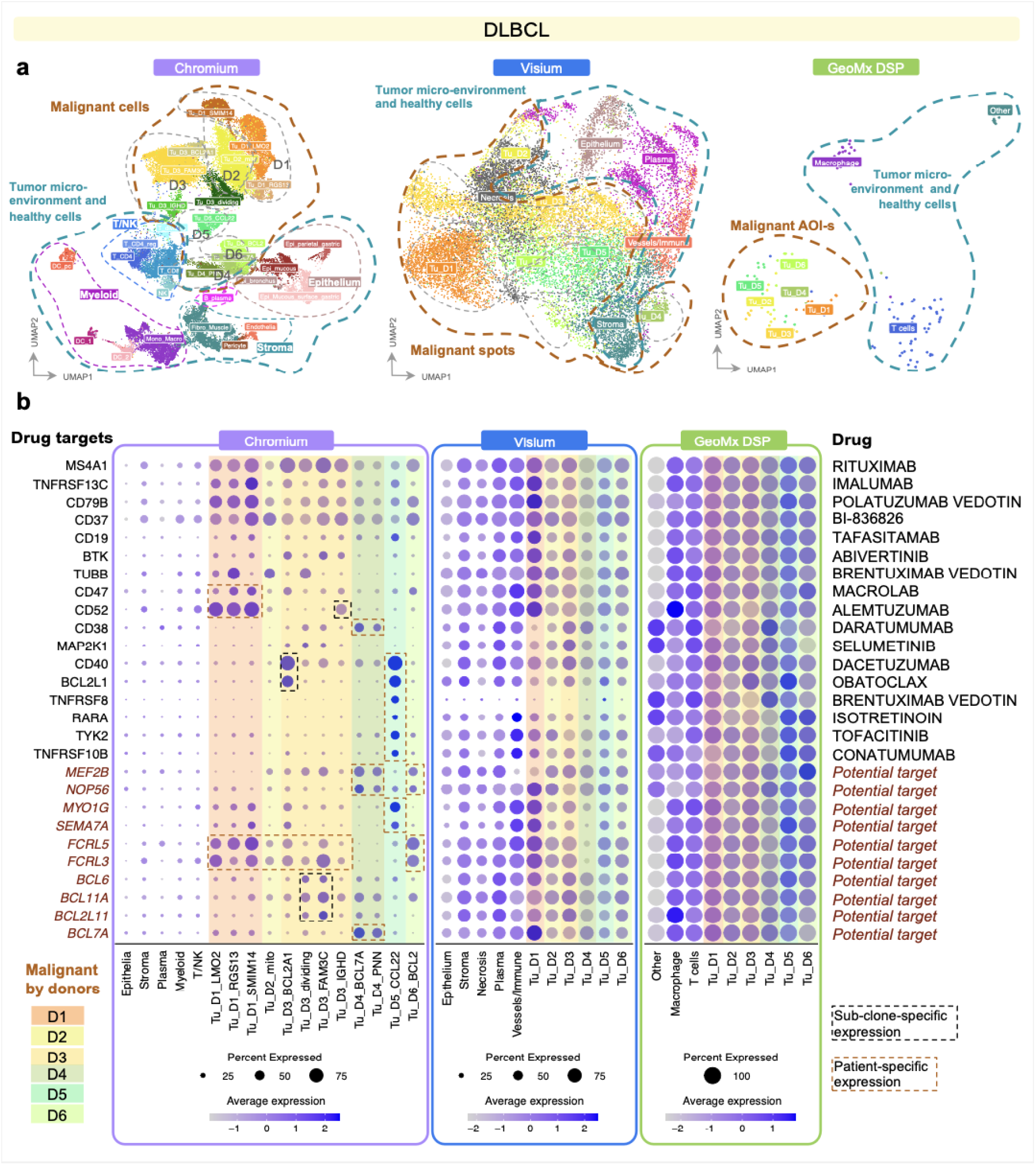
Exploration of drug targets and patient subgroups in DLBCL. (a) UMAP representations of Chromium, Visium and GeoMx DLBCL patients, colored by Level 4 cell types, clustering annotation of spots, or AOI types correspondingly. (b) Expression of selected genes that are significantly differentially expressed in at least one donor subtype compared to TME cells in any of the three methods. The drugs for known drug targets are listed per row. Genes with no drug are labeled as “Potential target”. Gene expression in any of the donors is color-coded. Patient-specific and subclone specific gene expression is highlighted with dashed lines.

Known drug targets were selected for exploring intra- and interpatient variability (Figure 6b). The B-cell markers were enriched in all patients’ malignant cells compared to the TME cells with Chromium data. We observed differences in expression between patients for many drug targets including CD47, CD52, CD40 and CD38, potentially suggesting an existing patient subgroup that might better respond to antibody-drug conjugates (ADC) designed against those surface molecules. Interestingly, we also observed intra-patient variability in expression in some of the drug targets. CD52 and some BCL genes showed varying expression within patient D3 between the identified subclones in single-nuclei data. Other potential drug targets such as FCRL genes showed patient-specific expression in D1, D2, D3 and D6. Interestingly, patient D5 stood out with very specific gene expression patterns, suggesting more drastic differences in the underlying biology.

While there was considerable heterogeneity both within and between patients’ malignant nuclei in the Chromium data, most of these differences were not observed in the Visium or GeoMx methods, likely due to the mixing of cells within spots or AOI-s. We tried to enhance the cell type signal specificity in Visium and GeoMx, by using deconvolution information with Chromium. Instead of the clustering approach above, we used majority votes from deconvolution, and showed densities of each cell type fraction for its corresponding majority vote class (Supplementary Figure 9a, b). To enhance the purity of spots, only spots with more than 50% maximum cell type fraction were kept in Visium (Supplementary Figure 9c). After such filtering, 52% DLBCL spots were included in the downstream analysis for Visium. For GeoMx, we kept 75% segments that showed consensus between AOI label and deconvolution majority vote label (Supplementary Figure 9d). The “purified” Visium and GeoMx data revealed a more distinct pattern in the expression levels of known drug targets, more similar to the patterns observed in Chromium, particularly for patient D5 (Supplementary Figure 9e). However, a lack of cell type specific gene expression persists in both Visium and GeoMx.

This use case is prominent in DLBCL where cellular density in tissue is high and heterogeneous, demonstrated by higher additional signals compared to breast and lung (Figure 2a, b). For such tissues with mixtures of cell types, Chromium provides the best resolution, while still missing the spatial coordinates. Overall, we demonstrate the ability to capture intra- and inter-patient variability in drug target expression with Chromium, but not Visium and GeoMx in DLBCL. This further emphasizes the importance of incorporating matched single-cell/nuclei RNA-seq data alongside full transcriptome spatial data to fully capture the complexity of cellular heterogeneity.

## Discussion

We aimed to compare full transcriptome methods on archival FFPE blocks from tumor patient samples: GeoMx DSP, Visium CytAssist and Chromium Flex. We assessed the operational challenges, specificity of cell type signatures, ability to capture heterogeneity and potential for high-throughput set-up.

The operational setup of GeoMx required more resources in terms of optimization, cost, and lab personnel training compared to Visium and Chromium. However, it offered greater flexibility in experimental design, despite being more susceptible to batch effects (Table 1). Chromium had about twice lower running cost, making the user consider whether spatial resolution on cell type groups is twice more valuable than corresponding single-cell/nuclei resolution. However, Chromium is suggested to consume at least five times more tissue than the spatial methods^3^, making this an important consideration for low-input samples such as core-needle biopsies. Estimated maximum throughput per one machine is highest for Chromium and lowest for GeoMx with the current experimental set-up. Pre-processing and analysis of the data is possible with user-provided tools, but advanced analyses are easily available for Visium and Chromium with a large scientific community and contributors but require more specialized effort for GeoMx.

**Table 1.**
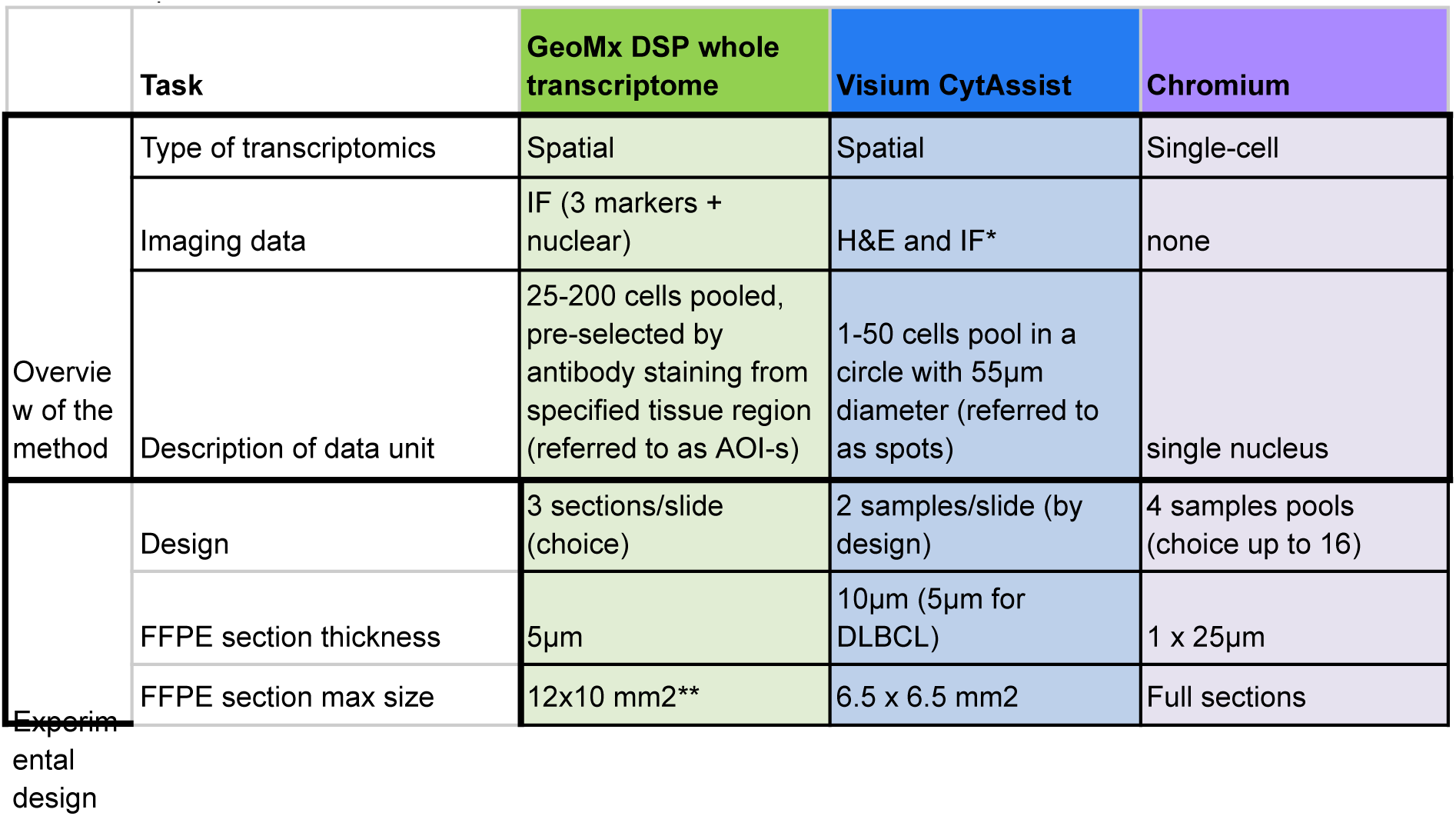
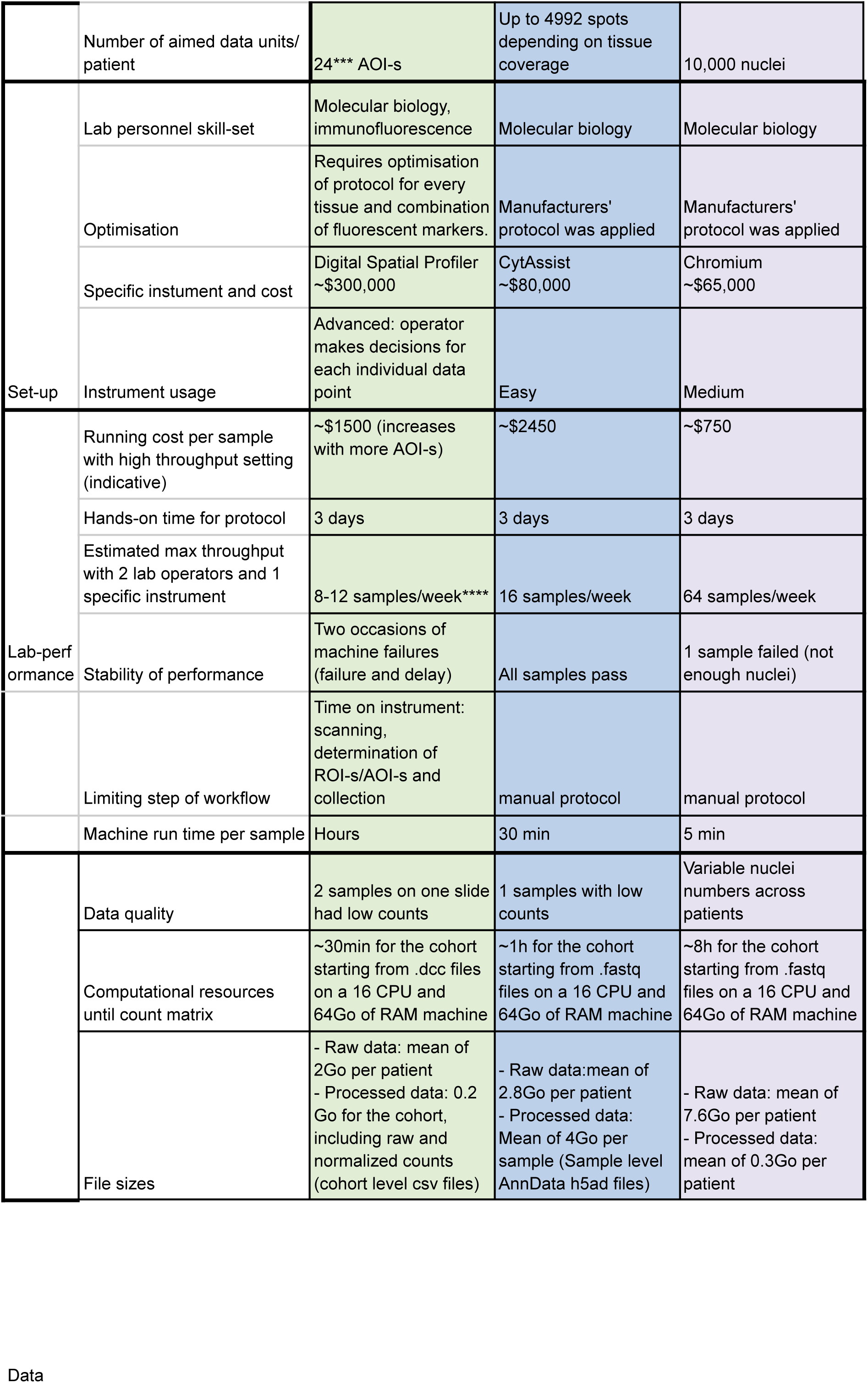
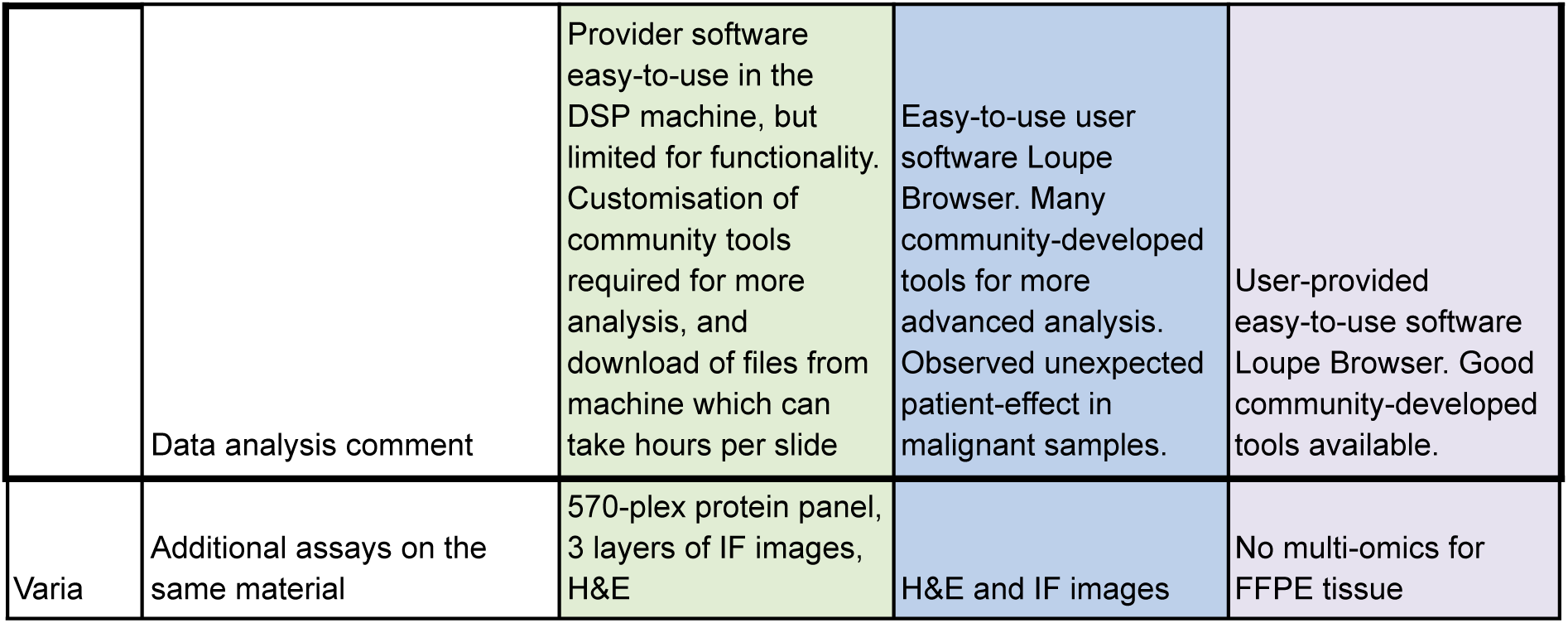
Comparison of the methods.

Although the data was highly reproducible across similar biological samples for all three methods, GeoMx and Visium exhibited greater biological variability across samples due to the mixing of cells within individual data points. This variability is inherent to Visium, as its capture of transcripts within circular spots on the tissue naturally leads to cell mixtures, which was expected. However, it was surprising to observe high fractions of other cell types in the GeoMx AOI-s, which were preselected for specific cell types — defeating the purpose of targeted selection to some extent. This issue was particularly pronounced in scattered cell types within the tissue, potentially due to transcript or probe leakage from nearby areas or physically overlapping cells along the Z-axis of the tissue. Contamination in GeoMx could be reduced with a simpler experimental design, such as selecting fewer segments or better-separated cell clusters, but making the flexibility of GeoMx a less relevant advantage of the method.

While the challenges of setup and unexpected transcript signals in GeoMx could have been mitigated with a simpler design, we aimed for the highest resolution of cell types, as this was the biggest advantage of GeoMx over Visium. Indeed, we collected CD68-enriched data from GeoMx that were not matched with Visium, highlighting both the benefit and bias of the highly flexible GeoMx method. The current best full-transcriptome resolution would be the Visium HD method with a 2×2μm grid^19^. However, this method is much more expensive, making it less applicable to larger studies. The highest spatial resolution would be provided by in situ sub-cellular CosMx and Xenium methods that can detect up to 6000 genes^20^ and more recently even full-transcriptome^21^ However, the costs for these in-situ technologies are even higher, and their throughput is at multiple days per sample, not allowing scale-up for study across a large number of patients. The development of sub-cellular technologies into full-transcriptome assays is the next frontier in studying tissue biology, with the cost and analysis difficulty hindering the usage by scientific and clinical communities.

While GeoMx can provide enrichment of chosen cell type transcriptomes, we demonstrated capture of tumor heterogeneity with both Visium and Chromium due to the nature of unbiased profiling across the tissue. However, the limitations of the original sample itself need be considered. In case of large tumors, multiple samples may be needed to describe all existing cell subsets.

Finally, we explored the functionality of full transcriptome methods in drug target discovery from highly heterogeneous FFPE tissue. Expression of known and novel drug targets in DLBCL varied across patients and within patients in with Chromium data, showing promise for patient subgroup identification and for assigning personalized therapies. Expression differences were however not clear in Visium and GeoMx, probably due to the cell type mixtures in each data unit. While all three methods have the potential for the discovery of targets, GeoMx is limited in the number of data points for intra-patient heterogeneity discovery, and both GeoMx and Visium are limited in cell type specific signal.

Overall, we successfully applied full-transcriptome methods in tumor FFPE blocks and demonstrated the applicability of Visium and Chromium for the systematic spatial profiling of large panels of samples. The unbiased nature of these platforms make them ideal in high throughput setups aiming at characterizing thousands of tumor samples. Based on this data, we propose Visium and Chromium as the platform of choice for spatial and transcriptomic analysis of tumor samples as part of the MOSAIC (Multi-Omics Spatial Atlas in Cancer) project^22^. The MOSAIC consortium will systematically profile thousands of tumor samples to analyze their spatial and single cell structures. By integrating these data with other modalities, like H&E staining, using advanced AI, we aim to create meaningful representations of cancer histology and transcriptomics. Coupled with clinical outcomes, these insights will offer an unprecedented understanding of how tumor spatial organization influences disease progression and treatment response.

## Methods

### Human tissue samples

The institutional review board of the Lausanne University Hospital and the local ethics committee approved the study, in accordance with the Helsinki Declaration (CER-VD 2023-00080). Written informed consent was available for all patients. Tissue sections were obtained from archival FFPE tumor samples. Histological diagnoses were reviewed by consensus between two pathologists (KvL and CS for Breast and Lung; LdL and CS for DLBCL). Detailed information on histotype, block age, and DV200 values are listed in Supplementary Table 1.

### Chromium FLEX

#### Lab workflow

For the single-cell transcriptomics, two FFPE curls of 25 micrometers were sectioned from each of the FFPE block. Nuclei were extracted using snPATHO protocol^9^, based on the protocol from 10X (CG000632). The nuclei were counted on the LUNA-FX7 cell counter (AO/PI viability kit, Logos). Pools of four samples were processed together as shown in Supplementary Table 4, targeting 40,000 nuclei recovery per pool (10,000 nuclei per sample). Replicas for two donors B3 and B4 were processed independently. The protocol was followed by the manufacturer’s protocol Chromium Fixed RNA profiling - Multiplexed samples (CG000527). Libraries were sequenced using a NovaSeq™ 6000/X sequencer aiming for 10,000 read pairs/nuclei. bcl2fastq was used for demultiplexing libraries after sequencing. FASTQ files were processed with Cell Ranger 7.1.0 (10X Genomics) with multi pipeline and human genome reference GRCh38-2020-A.

#### Pre-processing, QC (quality control) and annotation

Filtered barcode counts matrices from CellRanger were imported into R (v. 4.3.2, https://www.R-project.org/) analyses by the Seurat package (v. 5.0.3) ^23^. SoupX ^24^ was used to remove ambient RNA contamination. Only cells with minimum 200 genes (and max 6000 genes in DLBCL only), and <10% (and <20% in breast and lung) of reads mapping to mitochondrial genes were kept. For each sample, corrected raw counts were normalized using SCTransform. The top 3000 variable genes across samples were selected using the SelectIntegrationFeatures function. Dimensionality reduction using principal component analysis (PCA) was done, followed by a Uniform Manifold Approximation and Projection (UMAP) dimensional reduction using 50 principal components. Clustering with shared nearest neighbor (SNN) modularity optimization-based clustering algorithm implemented in the FindNeighbors and FindClusters functions was performed, with 30 principal components and resolutions between 0.4 and 0.8. The expression level of canonical marker genes and the top differentially expressed genes were used for identifying known cell types corresponding to the clusters. We finally performed sub-clustering to increase the granularity of annotations

#### Deconvolution

Chromium data are used as reference for deconvolution with cell type annotation at different levels of resolution. Since breast and lung cancer samples are both solid tumor and are annotated together, when constructing Chromium reference, we combine healthy cell types from both indications and add indication-specific tumor cells at Level 4.

For Figure 2 and Figure 3 on deconvolution cell type specificity of GeoMx and Visium, we used a combination of Level 1 and Level 4 annotation so that we obtain T cells and Macrophage cell types of interest while keeping concise grouping of other healthy or tumor cell types. For breast and lung cancer, T cells cell type is composed of “T_CD4”, “T_CD8_exhausted”, “T_CTL”, “T_CXCL13”, “T_reg”, and “TNK_dividing” from Level 4 annotation. For DLBCL, “T_CD4”, “T_CD4_reg”, “T_CD8”, “T_dividing” are grouped as T cells. For breast and lung cancer, we grouped Macrophage and Monocyte from Level 4 annotation as Macrophage. Because in GeoMx, the marker used for staining the *Macrophage* region is CD68+, which is expressed in both macrophage and monocyte cell types. For DLBCL, “Mono_Macro” cell type is labeled as Macrophage. Myeloid else consists of “DC_1”, “DC_2”, “DC_activated”, “DC_pc”, “Granulocyte”, “Mast_cell” for breast and lung, and “DC_1”, “DC_2”, “DC_pc” for DLBCL.

For Figure 4 and Figure 5 on Visium enabling biology discovery, to show scatterpie plot of deconvolution result for each patient and to obtain the deconvolution cell type majority vote for each spot, the tumor signature in Chromium reference was filtered to each patient’s Level 4 tumor subtypes, if there are multiple subtypes.

#### Intra- and inter-patient tumor heterogeneity analysis

For Figure 6, our interest is to identify the expression level of drug targets in tumor cell types in Chromium. Therefore, we make a condensed aggregation of healthy cell types from Level 1 annotation, while separating out tumor subtypes within each patient, if there are multiple subtypes, by adopting Level 4.

In Supplementary Figure 9, to borrow insights from Chromium and enhance the drug discovery capability of Visium and GeoMx, healthy cell types at Level 1 and patient-specific tumor cells from Level 2 annotation are used for Chromium. For Visium, the density of deconvolution fraction for each targeted cell type in the corresponding majority voted cell type category were grouped from Level 4 to Level 1 for healthy spots and Level 2 for tumor spots. Similarly for GeoMx, the density of deconvolution fraction for each targeted cell type in the corresponding AOI labeled segments were combined in the following way: Cell type “Other” contains “Plasma”, “Epithelia”, “Fibro_muscle”, and “Vessel”; cell type “T cells” consists of “T-cell” and “NK”; cell type “Macrophage” is characterized as “Myeloid”.

Gene-drug matches in Figure 5, 6, and Supplementary Figure 9 were collected from the ChEMBL database^25^.

### GeoMx DSP

#### Lab workflow

All Breast, Lung and DLBCL samples were assayed on the Nanostring GeoMx Digital Spatial Profiler (DSP) platform using the Whole Transcriptome Atlas (WTA) probe panel with NGS readout. One to three tissue sections were placed on each slide (Supplementary Figure 1a). Two samples (one each for Breast and Lung, B1 and L1 respectively) were analyzed in duplicate. Due to the failure of the GeoMx DSP instrument, two samples were repeated, including L4 and a duplicate of L3.

Slides were processed according to the manufacturer’s instructions. Tissue sections were cut at 5 um thickness. They were baked at 60°C for 2 hours. Sections were stained on a BOND-RXm fully automated multiplexing immunohistochemical stainer (Leica Biosystems; Wetzlar, Germany) using the following immunofluorescent antibodies: for Breast and Lung, PanCK - AF532 (Clone AE1 + AE3 from Novus) diluted at 1/50, CD3 - AF647 (Clone CD3-12 from Biorad) diluted at 1/100, CD68 - AF594 (Clone KP1, from Santa Cruz) in concentration 0.25 ug/ml and SYTO 13 nuclear stain diluted at 1:10000 (Nanostring) in Buffer W for 1 h at room temperature; for DLBCL, CD20 - AF594 (Clone IGL/773 from BioTechne) diluted at 1/100, CD3 - AF647 (Clone CD3-12 from Biorad) diluted at 1/100, CD68 - AF488 (Clone KP1 from e-BioMed) at concentration 0.25 ug/ml and SYTO 83 nuclear stain diluted at 1:100000 (Invitrogen) in Buffer W for 1 h at room temperature.

Regions of interest (ROIs) were selected by pathologists, and segmented into areas of illumination (AOI-s) based on immunofluorescent markers (Supplementary Tables 3-4). The first segment was defined as double positive for CD3 and CD68 with the purpose of excluding autofluorescent structures (such as elastin fibers in lung) or debris, and it was not collected. For Lung and DLBCL samples, the *CD3-AOI-s* were collected first, followed by *CD68-AOI-s*. For Breast samples, *CD68-AOI-s* were collected first, followed by *CD3-AOI-s*. *PanCK-AOI-s* and *CD20-AOI-s* were collected third for solid cancers and DLBCL respectively. For five and four ROI-s in breast and lung replica samples respectively, only two segments of *PanCK+* and *PanCK-* were collected in that order, ignoring other markers. The remaining cells showing SYTO nuclear expression in the absence of any cytoplasmic marker were collected as the fourth marker-negative (*Other*) segment in BC. All the remaining surface of ROIs which had not been collected into any AOI was collected as marker-negative (*Other*) segments for Lung and DLBCL, independently of the presence of nuclei. With the exception of the Other segment of Lung and DLBCL, a segment was collected when it contained at least 50 cells. About 24 such AOI-s per patient were collected with AOI size varying from about 500µm^2^ to 300,000µm^2^ with an average size of 60,000µm^2^. Library preparation and sequencing was performed according to the manufacturer’s instructions and kits. Briefly, i5 and i7 indices were added, reactions pooled and purified, and libraries sequenced with paired-read sequences with 2 × 27 base pairs. Manual curation of the annotation file was performed to match the collected segments with a biological cell fragment as AOI label (Supplementary Table 3)

#### Pre-processing and QC (quality control)

The GeoMx NSG pipeline *GeomxTools*^26^ was used to convert FASTQ files into expression matrices of raw probe counts stored in DCC files. AOI segment QC was conducted using NanoString recommendations: segments with >1000 raw reads, < 80% aligned, trimmed or stitched, < 50% saturation, >1000 NTC and < 100 nuclei are removed. There were 2 breast segments removed due to low saturation, 1 lung segment excluded due to low alignment, and 7 DLBCL segments were filtered out due to low reads (n = 1), low saturation (n = 2), and small nuclei area (n = 4). This results in a sample size of 122, 117, and 137 for breast, lung, and DLBCL samples, respectively.

Probes with geometric mean from all segments divided by the geometric mean of all probe counts representing the target from all segments less than 0.1, together with probes flagged as outliers according to the Grubb’s test in at least 20% of the segments, were filtered out. Gene raw counts were generated using the geometric mean of the associated probe counts. Meta variable number of genes detected were derived and plotted in Figure 1d GeoMx panel.

Segments from each indication were combined by *GeomxTools* as a *SummarizedExperiment* object, and then converted by *standR* ^27^ to a *SpatialExperiment* object. No AOI segment or gene was further excluded with the *standR* preprocessing pipeline.

#### Normalization and batch correction

As suggested by van Hijfte et al.^28^, we use quantile normalization for GeoMx data analysis. Quantile normalization (*preprocessCore::normalize.quantile()*) was applied to *log1p()* transformed data. The reduced dimension of normalized GeoMx breast samples is shown in Figure 1d. We applied batch correction method RUV4, as implemented in *standR,* to quantile normalized data to remove batch effects introduced by slides. Distinct *T cells* and *Macrophage* clusters on the reduced dimension of normalized and batch corrected data are shown in Supplementary Figure 7a.

#### Deconvolution

Cell type abundance was estimated using the *SpatialDecon* package ^10^ and the signature matrix derived from the Chromium single-cell RNAseq dataset. When running deconvolution, we exponentiated back the log transformed, quantile normalized, and RUV4 batch corrected assay to obtain normalized and batch corrected data on linear scale, as suggested by *SpatialDecon::spatialdecon()* manual. To enhance the drug discovery potential of GeoMx, we borrowed insights from Chromium through deconvolution as shown in Supplementary Figure 9b, d, e.

### Visium CytAssist

#### Lab workflow

Sections with 10 micrometers for lung and breast, and 5 micrometers for DLBCL were placed on individual positively charged slides (Superfrost Plus Adhesion Microscope Slides, thermofisher). The slide preparation workflow was performed using the Visium CytAssist Spatial Gene Expression for FFPE by 10X (CG000520 and CG000518). High resolution H&E imaging was done on Aperio CS at 40x magnification. The 6.5mm x 6.5 mm capture area was chosen by the pathologists. Manufacturer’s workflow was followed until the library preparation. Libraries were sequenced with paired-end dual-indexing (28 cycles Read 1, 10 cycles i7, 10 cycles i5, 90 cycles Read 2) on NovaSeq™ 6000/X. Space Ranger 2.0.1 pipeline was used to align the reads to human genome reference GRCh38-2020-A. High resolution image and CytAssist image were aligned using Space Ranger.

#### Manual annotation of the Visium spots

Loupe Browser version 6.5.0 was used by the pathologists to annotate all of the Visium spots in one of the following classes: pure tumor (invasive carcinoma), in situ carcinoma, tumor-stroma mix, intratumoral stroma, lymphocytes, immune cell mix, tumor infiltrating lymphocytes, intratumoral vessels, artefact/ fold/ empty (for spots to be excluded from the analysis due to sectioning artefacts or falling outside the tissue), most likely tumor (for epithelial proliferations which were difficult to classify based on H&E staining alone), acellular mucin, necrosis/debris for solid Breast and Lung cancer; tumor, small lymphocytes, stroma, necrosis, epithelium, and vessels for DLBCL. Visium spots, each covering a mixture of cells, were assigned to one of the previously listed classes according to the majority cell type (> 50%) observed on H&E staining. The hybrid category “tumor-stroma mix” was used for spots covering an approximately equivalent mixture of tumoral cells and stroma. Immune cell mix annotation class was used for spots covering a mixture of cells, including macrophages, lymphocytes, and plasmocytes (Supplementary Table 5, Supplementary Figure 1f).

#### Pre-processing and QC (quality control)

Using the filtered gene count matrix, we excluded spots that had fewer than 100 unique molecular identifiers (UMIs), more than 22% mitochondrial reads, or were located at the edge of the fiducial frame, unreasonably distant from the majority of the tissue. Additionally, spots identified by pathologists as artifacts, folds, or empty were also filtered out. We set a quality control threshold requiring genes to be detected in at least 20 spots to be included in the analysis.

#### Deconvolution

After QC, cell type fraction is estimated using deconvolution method *Cell2location*^11^, which takes raw unique molecular identifier (UMI) counts of each Visium sample and annotated single cell reference data as input. We defined a Visium annotation approach from deconvolution majority vote with Level 4 Chromium annotation, by setting a cut-off that if the highest cell type fraction is above 20%, we label the spot as that cell type, otherwise, we label the spot as “Mix”. To enhance the drug discovery potential of Visium, we borrowed insights from Chromium through deconvolution as shown in Supplementary Figure 9a, c, e.

#### Normalization

We adapted the recommended normalization method for different biological questions. To show sample reproducibility among replicates and to integrate samples within each indication, we followed the Seurat workflow ^23^ and used *SCTransform* normalization. The reduced dimension of normalized Visium breast samples is shown in Figure 1d Visium panel. On the other hand, we used the default log normalization for individual sample’s spatial aware clustering (Figure 5b) to achieve optimal spatial domain detection that detects intra-patient heterogeneity, as recommended in the BayesSpace vignette.

#### Clustering

For individual sample discovery (Figure 5b), we used the graph-based clustering approach implemented in Seurat to identify the optimal number of clusters for *BayesSpace* ^12^. We followed the default *BayesSpace* pipeline and parameters, with log normalization and 50 PCs. The log transformed subspot gene expression was inferred from subspot level PCs, with a linear model projection between spot level log transformed gene expression to spot level PCs, as detailed in *BayesSpace*. We used *Seurat::FindAllMarkers()* to find significantly (adjusted p value < 0.05) differentially expressed genes between clusters of interest. Gene set enrichment analysis was done with R package *enrichR* ^29^, and the gene set reference was from Molecular Signatures Database (MSigDB) Hallmark Gene Set Collection (2020) database.

#### Integration

To integrate multiple samples with the same indication across patients, we decontaminated Visium counts with SpotClean^30^ and used SCTransform normalization for each sample. *SpotClean::spotclean()* is run with the default parameter on the raw gene count matrix from the SpaceRanger output. We then performed QC of the spot-cleaned object by filtering to the gene and spot barcodes that passed QC previously. We used *Seurat::SCTransform()* in Seurat (v.5.0.3) ^23^ to normalize each spot-cleaned Visium sample and to identify variable genes. We used *Seurat::SelectIntegrationFeatures()* function to select a union of 3000 HVGs by consensus ranking of the gene from all samples. Standard Seurat pipeline (e.g. *RunPCA()*, *FindNeighbors()*, *FindClusters()*) is used to obtain the reduced dimension UMAP and clusters.

DLBCL Visium data was annotated into regions after Seurat clustering on the integrated object (Supplementary Figure 8a). Donor and pathology annotation distribution was considered for each cluster to annotate clusters 4 (D5) and 5 (D6) and clusters 2, 12, 14, 16 (necrosis) correspondingly (Supplementary Figure 8b, c). Canonical marker genes’ expression in each cluster (Supplementary Figure 8d) aided to annotate regions: clusters 9, 10 and 13 were mostly expressing plasma cell marker genes, cluster 6 was assigned to epithelium, and cluster 1 was showing strong stroma signal. Cluster 7 was likely a mixture of immune cells with vessels, which was confirmed by pathology annotation. Aggregated patient-specific tumor markers from Supplementary Figure 3 were used to annotate malignant areas: cluster 0 (D1), clusters 15, 18 (D2), clusters 3, 8, 11 (D3), and cluster 17 (D4) were identified. *Seurat::FindAllMarkers()* was further consulted for differentially expressed genes between clusters.

#### Visualization

Deconvolution cell type proportion across all spots in one sample, pathology annotation, clusters and other continuous values, such as gene expression and a single cell type deconvolution proportion, as well as reduced dimensions were visualized spatially with R packages *Seurat* ^23^ and *ggspavis* ^31^.

### Spot-segment matching between Visium and GeoMx

Fluorescent images carrying GeoMX’s regions of interest and areas of interest were registered to high resolution H&E images carrying Visium spots using the Elastix software^32,33^. For GeoMX images, registration was performed on the SYTO 13 channel, with pixel intensities clamped to 1% and 99% of extremal values in order to alleviate fluorescent artefacts. H&E images were converted to grayscale prior to registration. Registration was performed on downsampled images with a resolution of (approximately) 4 microns per pixel. Affine registration parameters were optimized by minimizing Mattes advanced mutual information metrics^32^. As GeoMX’s fluorescent image and Visium’s H&E were carried on two subsequent slides of the same block, a visual inspection was performed to ensure the quality of the registration procedure. For mapped spots and subspots, PanCK- AOI-s are excluded from the analysis due to their impurity by definition.

## Data and code availability

All the data files are available at ArrayExpress database (http://www.ebi.ac.uk/arrayexpress) under accession numbers E-MTAB-14560 and E-MTAB-14566. Single-cell data is available for browsing at cellxgene database: https://cellxgene.cziscience.com/collections/bd552f76-1f1b-43a3-b9ee-0aace57e90d6. Data analysis code is available in https://github.com/bdsc-tds/mosaic_pilot_study.

## Legends

### Supplementary Figures

Supplementary Figure 1: Overview of data

(a) Schematic for GeoMx and Visium spatial transcriptomics and Chromium scRNA-seq data generation for breast, lung and DLBCL cancer types. (b) Total number of transcripts per AOI, spot, and cell in GeoMx, Visium, and Chromium, respectively, across all samples and indications. (c-e) Sample size with number of nuclei, spots and AOI-s in Chromium, Visium and GeoMx, respectively, for all indications. (f) Percentage of spots in Visium falling into each pathology annotation for indications. (g) Number and percentage of AOI-s in GeoMx falling into each curated segment as AOI label in all indications. AOI - area of illumination. DLBCL - Diffuse Large B-cell Lymphoma. DSP - digital spatial profiler.

Supplementary Figure 2: Chromium scRNA-seq data annotated into different levels of cell type groups

Breast and Lung (a) and DLBCL (b) samples received level 4 annotation based on unsupervised clustering and canonical marker expression. The higher levels were derived by combining cell types of the lower level into biologically meaningful groups.

Supplementary Figure 3: Cell type frequencies and malignant marker genes on level 1 and level 4 annotated Chromium

(a) Ordered by category percentage, the cell type frequency in each indication at Level 1 annotation. Ordered by category percentage, the marker genes for malignant nuclei in each donor, and cell type frequency at Level 4, ordered by percentage for breast (b), lung (c) and DLBCL (d).

Supplementary Figure 4: Cell type mixtures in GeoMx segments by marker gene expression and unsupervised clustering

Cell type specific genes’ expression in AOI-s grouped by collected AOI label in breast and lung samples (a) and in DLBCL (b). Known marker genes’ expression specific for the selected segment is highlighted in light purple as “Expected” signal. “Unexpected” signal shows expression of the known marker in other segments. (c) Heatmap of normalized and batch corrected gene expression matrix on top 2000 differentially expressed genes between AOI label types, annotated by unsupervised clustering and metadata from Lab Worksheet of GeoMx breast samples (Supplementary Table 4b). Highlighted areas corresponding to AOI labels.

Supplementary Figure 5: A gallery of breast and lung Visium samples with pathology annotation, deconvolution majority vote, and GeoMx matching

(a) Sample names matched between GeoMx and Visium for registration analysis as consecutive sections from the same block in breast and lung. Breast and lung Visium samples colored by pathology annotation (b) and deconvolution majority vote (c). For samples that are used for registration, we plot spots (d) that are mapped to an AOI label spatially, colored by their mapped AOI label.

Supplementary Figure 6: A gallery of DLBCL Visium samples with pathology annotation, deconvolution majority vote, and comparison of their agreement in each indication.

DLBCL Visium samples colored by pathology annotation label (a) and Level 4 deconvolution majority vote (b). (c) Agreement between pathology annotation and cell type deconvolution result, displayed as average cell type deconvolution fraction per pathology label per cell type, for breast, lung, and DLBCL samples, separately. Expected signal is surrounded by a green box.

Supplementary Figure 7: Immune cell signals on reduced dimensions of integrated Visium and GeoMx, and intra-patient heterogeneity discovery with Chromium and Visium.

(a) UMAP plots in Visium and (b) TSNE plots in GeoMx integrated between all donors for each indication, showing enrichment of T-cell and Macrophage cell type groups by deconvolution fractions. *T cells* and *Macrophage* AOI labels on GeoMx are shown by triangles. (c) Different deconvolution fractions of tumor and fibroblast cell types between Area A and B. (d) UMAP of Chromium B3 tumor nuclei with clusters gained by unsupervised clustering, annotated into Tu_B3_PLA2G2A and Tu_B3_NPPC malignant cell substates. Marker genes PLA2G2A and NPCC expression on UMAP. (e) Spatial localisation of the two tumor subclusters to Areas A and B on patient B3 by deconvolution cell type fraction. (f) DE of the two tumor substates. (g) Gene set enrichment of DE genes from (f).

Supplementary Figure 8: Annotation of DLBCL Visium spots into regions.

(a) UMAP of integrated DLBCL Visium samples, colored by Seurat cluster, with final annotations into regions. (b) Heatmap showing proportion of each cluster that belongs to spots from which patient. Due to the imbalance of sample size (number of spots) in DLBCL Visium samples, we weighted the counts in each sample by the inverse of its sample size. (c) Heatmap showing proportion of each cluster that belongs to spots of which pathology annotation class. (d) Clustered dot plot showing canonical markers of known healthy cell types and gene expression level across all clusters, highlighting the assigned clusters to regions. (e) Aggregated patient-specific tumor marker (based on patient-specific malignant cell markers in Chromium in Supplementary Figure 3b-d) gene expression on UMAP of integrated DLBCL Visium samples.

Supplementary Figure 9: Chromium informed data-driven Visium and GeoMx inter-patient drug discovery.

Deconvolution fractions in DLBCL for each targeted cell type in the corresponding majority voted cell type category in Visium (a) and for each targeted cell type in the corresponding AOI labeled segments in GeoMx (c). Higher purity is achieved by restricting the analysis to spots where the deconvolution majority cell type label’s fraction is > 50% in Visium (b) and by filtering for segments where the AOI label and the deconvolution majority vote are in consensus in GeoMx (d). (e) Expression of drug targets as in Figure 6 with improved purity spots and AOI-s in Visium and GeoMx.

### Supplementary Tables

Supplementary Table 1: Patient overview

A table with patient ID, Sample type, Cancer type, Tissue, Histology type, histology specification for Invasive lobular carcinoma, Immune histology, DV200 (%) as RNA quality value, Block age in months. NOS - Not Otherwise Specified. *Patient had a replica sample for GeoMx and Visium, **Patient had a replica sample for Visium, ***Patient had a replica sample for GeoMx.

Supplementary Table 2. Single-cell samples

Information about single-cell samples: Sample ID, Patient ID, Sample repeating info, Cancer type, Chromium pool ID, Number and thickness of FFPE curls (micrometers), Pass/Fail status.

Supplementary Table 3. GeoMx samples

Information about GeoMx samples: GeoMx slide ID, GeoMx run date, Patient ID, Section ID, Section letter on slide (A - closest to the label, C - furthest from the label if not otherwise specified), Comment on Pass/Fail status, FFPE section thickness, Staining mix with morphology marker information, Number of AOI-s collected. Calculated for each indiation in µm^2^: Minimum AOI area, Maximum AOI area, Mean AOI area, SD AOI area. AOI - area of illumination. SD - standard deviation.

Supplementary Table 4a. DLBCL Lab Worksheet

Information about GeoMx AOI-s in DLBCL. Sample ID as AOI unique ID used in sequencing files, GeoMx slide name, GeoMx DSP scan name, transcriptome panel, Region of Interest ID (roi), Segment type based on morphology markers, Area of Illumination type (aoi), AOI area size in µm^2^, Patient ID, Manually curated Cell fraction based on immunofluorescence image of morphology markers and the GeoMx DSP mask, AOI label with Manual curation, final AOI label with categories.

Supplementary Table 4b. Breast Lab Worksheet

Information about GeoMx AOI-s in Breast as in DLBCL lab worksheet. Additional values: ROI name, ROI type, ROI_category, AOI label with location, Indication, Patient ID, Section ID, Unique ID human readable (hr), Comment, Tags, number of nuclei in AOI, ROI Coordinate X, ROI Coordinate Y, Scan Date.

Supplementary Table 4c. Lung Lab Worksheet

Information about GeoMx AOI-s in Lung. All the same values as for Breast.

Supplementary Table 5. Pathology annotation distribution.

Pathology annotations, their grouping categories for Figure 2, suggested colors, and number of spots in each sample with the annotation.

Supplementary Table 6. Single-cell annotations in DLBCL

Single-cell annotations at different levels, suggested colors and ontology terms. Level4 annotations, Harmonised Level 4 annotations across lung, breast and DLBCL, Harmonized colors suggestions, cell_type_ontology_term_id: CL, all other labels for Levels 3, 2.5, 2, 1.5 and 1. Level 1.5 generated as a mixture from Level 1 to 4 with cell type groups most similarly matching GeoMx segments. Matching equivalent GeoMX segment and colors.

Supplementary Table 7. Single-cell annotations in Breast and Lung

Single-cell annotations at different levels, suggested colors and ontology terms as in Supplementary Table 6.

## Citations

1. Method of the Year 2020: spatially resolved transcriptomics. Nat. Methods 18, 1 (2021).

2. Du, M. R. M., et al. Spotlight on 10x Visium: a multi-sample protocol comparison of spatial technologies. bioRxiv 2024.03.13.584910 (2024) doi:10.1101/2024.03.13.584910.

3. Janesick, A. et al. High resolution mapping of the tumor microenvironment using integrated single-cell, spatial and in situ analysis. Nat. Commun. 14, 8353 (2023).

4. Zimmerman, S. M. et al. Spatially resolved whole transcriptome profiling in human and mouse tissue using Digital Spatial Profiling. Genome Res. 32, 1892 (2022).

5. Wu, H., Kirita, Y., Donnelly, E. L. & Humphreys, B. D. Advantages of Single-Nucleus over Single-Cell RNA Sequencing of Adult Kidney: Rare Cell Types and Novel Cell States Revealed in Fibrosis. J. Am. Soc. Nephrol. 30, 23 (2019).

6. Wang, T. et al. An experimental comparison of the Digital Spatial Profiling and Visium spatial transcriptomics technologies for cancer research. bioRxiv 2023.04.06.535805 (2023) doi:10.1101/2023.04.06.535805.

7. Cervilla, S. et al. Comparison of spatial transcriptomics technologies across six cancer types. bioRxiv 2024.05.21.593407 (2024) doi:10.1101/2024.05.21.593407.

8. Andrews, T. S. et al. Single-cell, single-nucleus, and spatial transcriptomics characterization of the immunological landscape in the healthy and PSC human liver. J. Hepatol. 80, 730–743 (2024).

9. Vallejo, A. F., et al. snPATHO-seq: unlocking the FFPE archives for single nucleus RNA profiling. bioRxiv 2022.08.23.505054 (2022) doi:10.1101/2022.08.23.505054.

10. Danaher, P. et al. Advances in mixed cell deconvolution enable quantification of cell types in spatial transcriptomic data. Nat. Commun. 13, 385 (2022).

11. Kleshchevnikov, V. et al. Cell2location maps fine-grained cell types in spatial transcriptomics. Nat. Biotechnol. 40, 661–671 (2022).

12. Zhao, E. et al. Spatial transcriptomics at subspot resolution with BayesSpace. Nat. Biotechnol. 39, 1375–1384 (2021).

13. Castellano, E. & Downward, J. RAS Interaction with PI3K: More Than Just Another Effector Pathway. Genes Cancer 2, 261–274 (2011).

14. Yu, H., Pardoll, D. & Jove, R. STATs in cancer inflammation and immunity: a leading role for STAT3. Nat. Rev. Cancer 9, 798–809 (2009).

15. Laplante, M. & Sabatini, D. M. mTOR signaling in growth control and disease. Cell 149, 274–293 (2012).

16. Buck, M. B. & Knabbe, C. TGF-beta signaling in breast cancer. Ann. N. Y. Acad. Sci. 1089, 119–126 (2006).

17. Wang, Y. & Zhou, B. P. Epithelial-mesenchymal transition in breast cancer progression and metastasis. Chin. J. Cancer 30, 603–611 (2011).

18. Bidwell, B. N. et al. Silencing of Irf7 pathways in breast cancer cells promotes bone metastasis through immune escape. Nat. Med. 18, 1224–1231 (2012).

19. Oliveira, M. F. et al. Characterization of immune cell populations in the tumor microenvironment of colorectal cancer using high definition spatial profiling. bioRxiv 2024.06.04.597233 (2024) doi:10.1101/2024.06.04.597233.

20. CosMx^TM^ Human 6K Discovery Panel. NanoString https://nanostring.com/products/cosmx-spatial-molecular-imager/cosmx-rna-assays/hum an-6k-discovery-panel/ (2023).

21. CosMx^TM^ human whole transcriptome panel. NanoString https://nanostring.com/products/cosmx-spatial-molecular-imager/cosmx-rna-assays/human-whole-transcriptome-panel/ (2024).

22. MOSAIC is the world’s largest spatial multiomics dataset in oncology. https://www.mosaic-research.com/.

23. Hao, Y. et al. Dictionary learning for integrative, multimodal and scalable single-cell analysis. Nat. Biotechnol. 42, 293–304 (2024).

24. Young, M. D. & Behjati, S. SoupX removes ambient RNA contamination from droplet-based single-cell RNA sequencing data. Gigascience 9, (2020).

25. Zdrazil, B. et al. The ChEMBL Database in 2023: a drug discovery platform spanning multiple bioactivity data types and time periods. Nucleic Acids Res. 52, D1180–D1192 (2024).

26. GeomxTools. Bioconductor http://bioconductor.org/packages/GeomxTools/.

27. Liu, N. et al. standR: spatial transcriptomic analysis for GeoMx DSP data. Nucleic Acids Res. 52, e2 (2024).

28. van Hijfte, L. et al. Alternative normalization and analysis pipeline to address systematic bias in NanoString GeoMx Digital Spatial Profiling data. iScience 26, 105760 (2023).

29. Kuleshov, M. V. et al. Enrichr: a comprehensive gene set enrichment analysis web server 2016 update. Nucleic Acids Res. 44, W90–7 (2016).

30. Ni, Z. et al. SpotClean adjusts for spot swapping in spatial transcriptomics data. Nat. Commun. 13, 2971 (2022).

31. ggspavis. Bioconductor http://bioconductor.org/packages/ggspavis/.

32. Mattes, D., Haynor, D. R., Vesselle, H., Lewellyn, T. K. & Eubank, W. Nonrigid multimodality image registration. in Medical Imaging 2001: Image Processing vol. 4322 1609–1620 (SPIE, 2001).

33. Shamonin, D. P. et al. Fast parallel image registration on CPU and GPU for diagnostic classification of Alzheimer’s disease. Front. Neuroinform. 7, 50 (2013).

