## Supplementary figures and images for "Transcriptome Analysis of Archived Tumor Tissues by Visium, GeoMx DSP, and Chromium Methods Reveals Inter- and Intra-Patient Heterogeneity"

### Supplementary figures

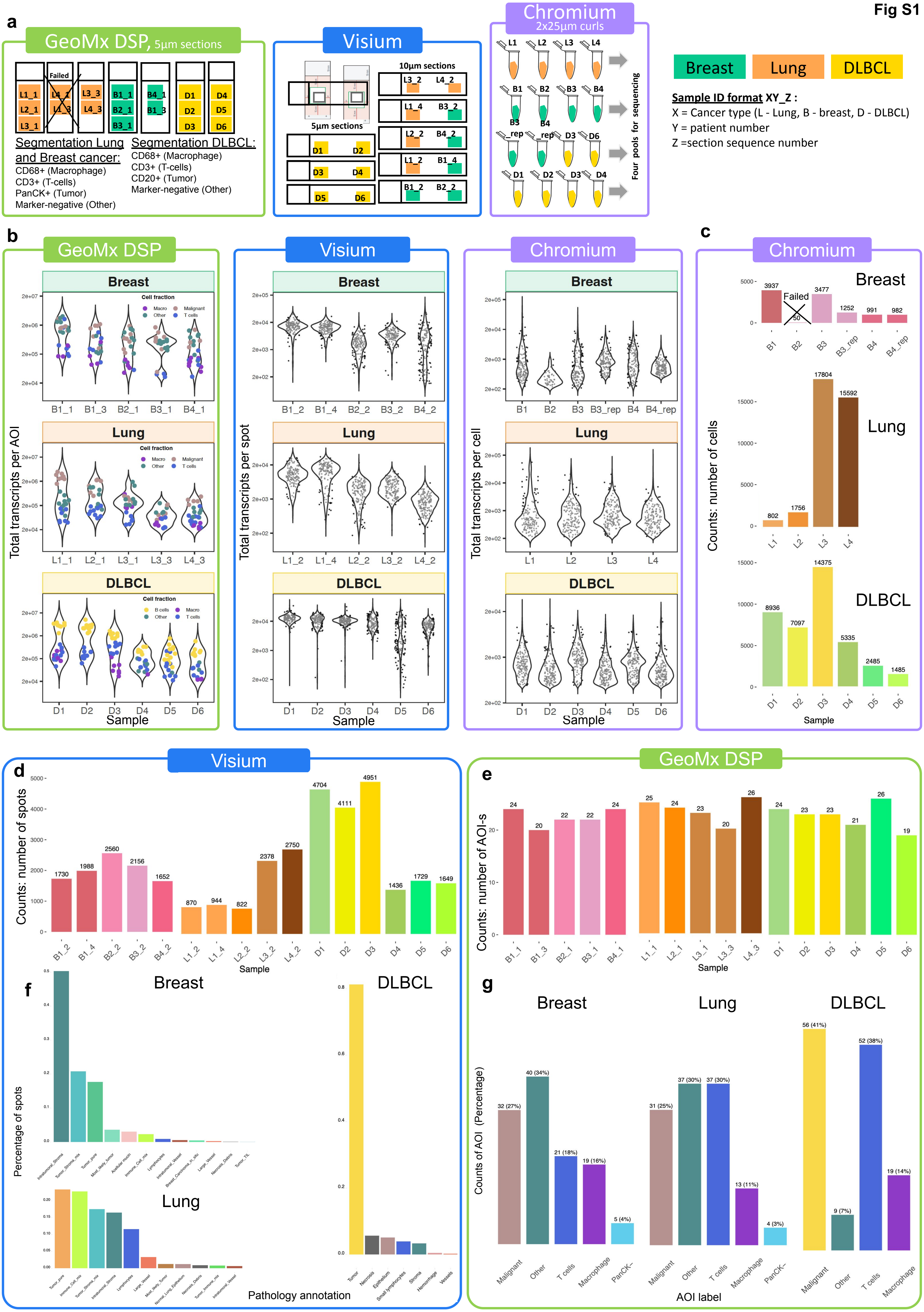

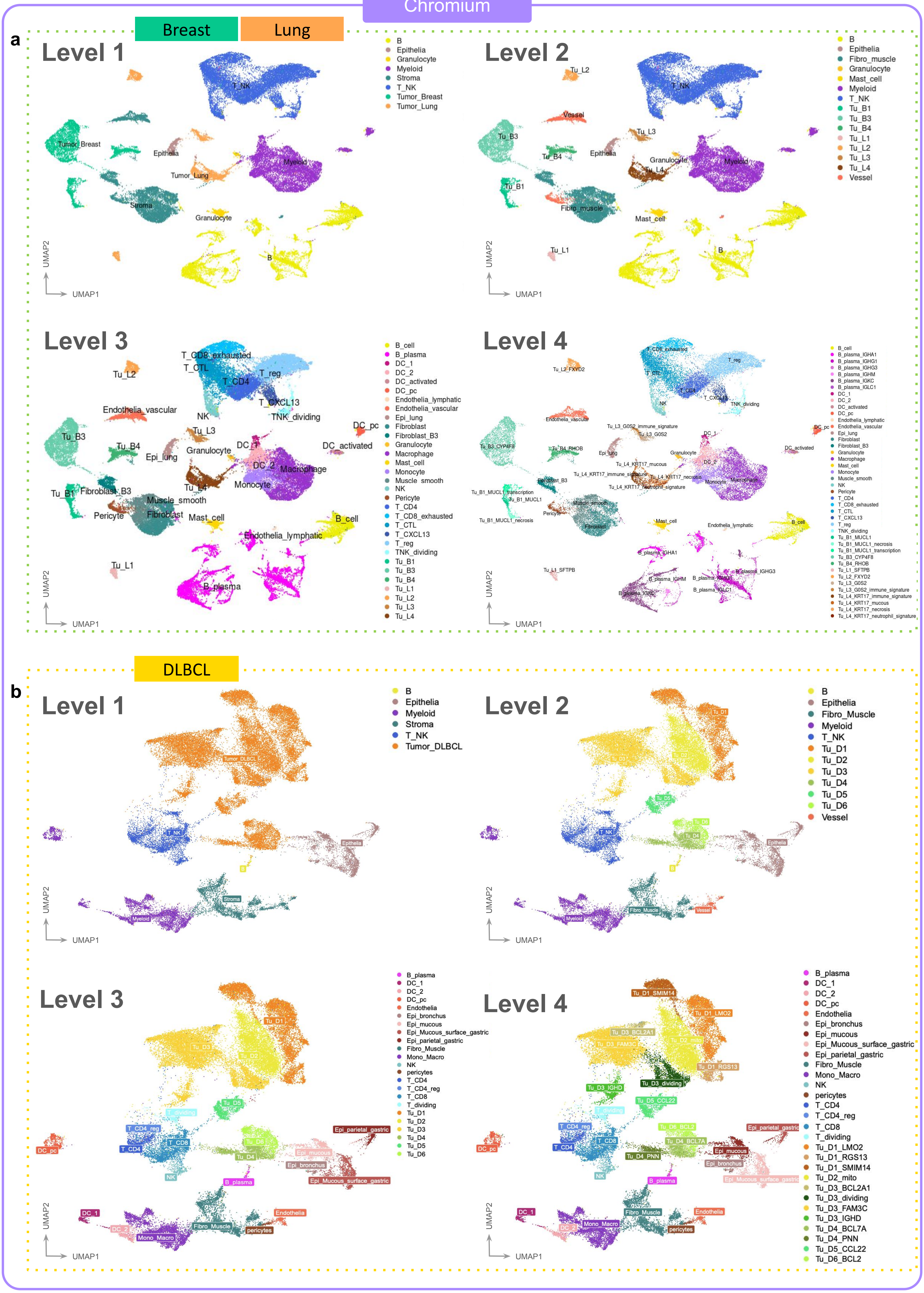

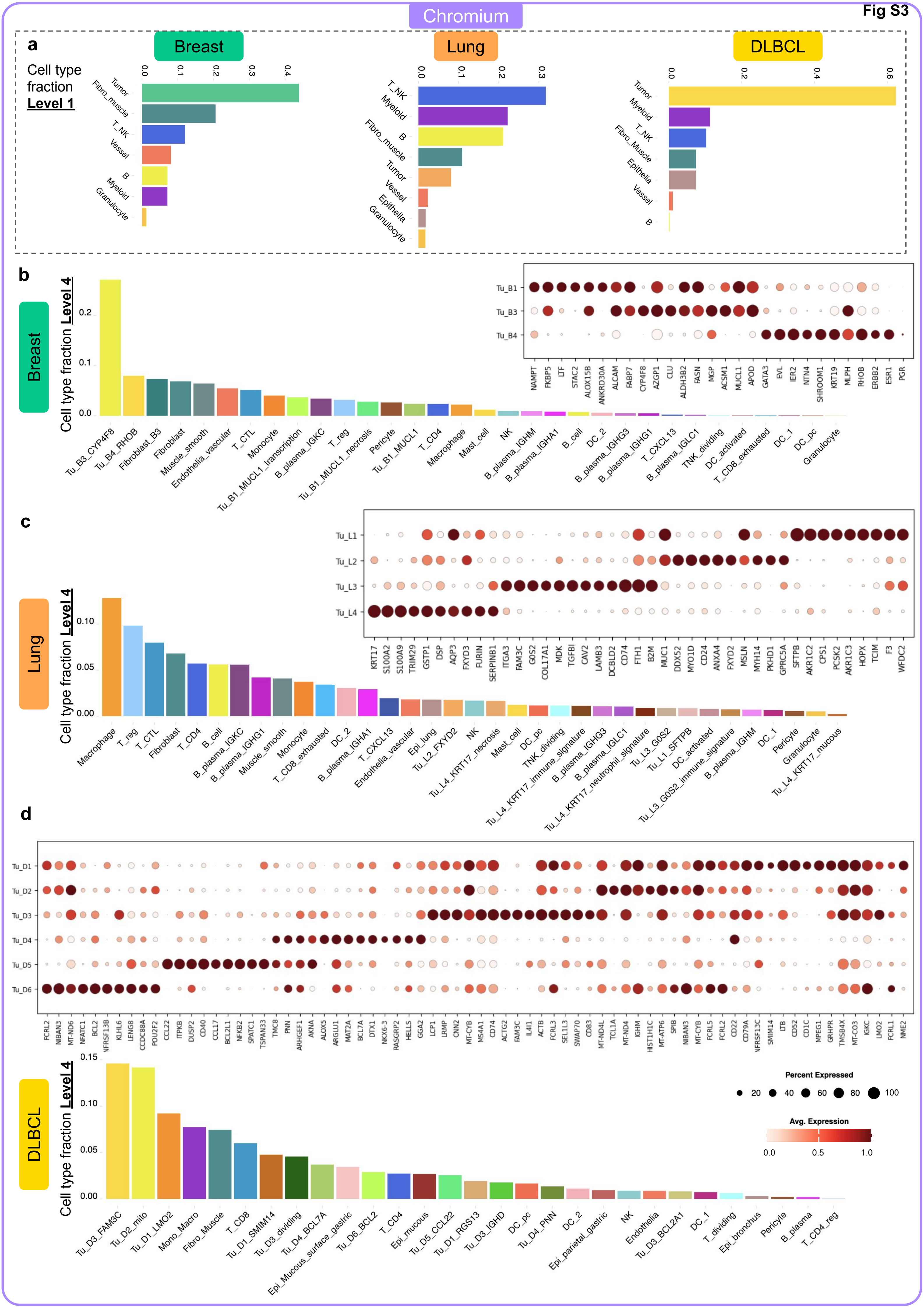

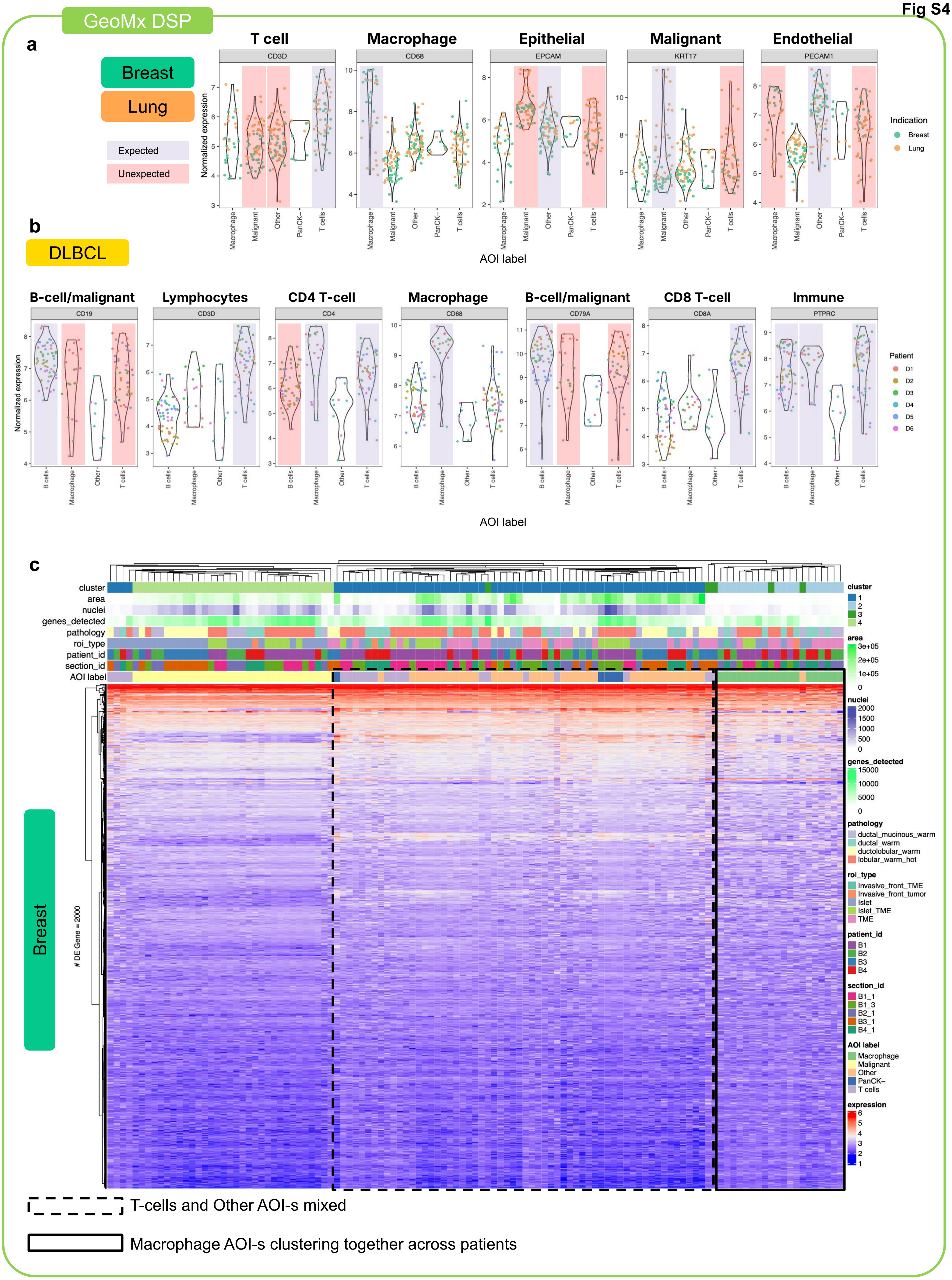

Fig S5

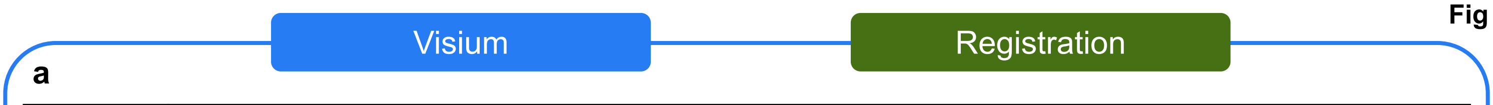


| GeoMx | B1_1 | B1_3 | B2_1 | B3_1 | B4_1 | L1_1 | L2_1 | L3_3 | L4_3 |
|---|---|---|---|---|---|---|---|---|---|
| Visium | B1_2 | B1_4 | B2_2 | B3_2 | B4_2 | L1_2 | L2_2 | L3_2 | L4_2 |

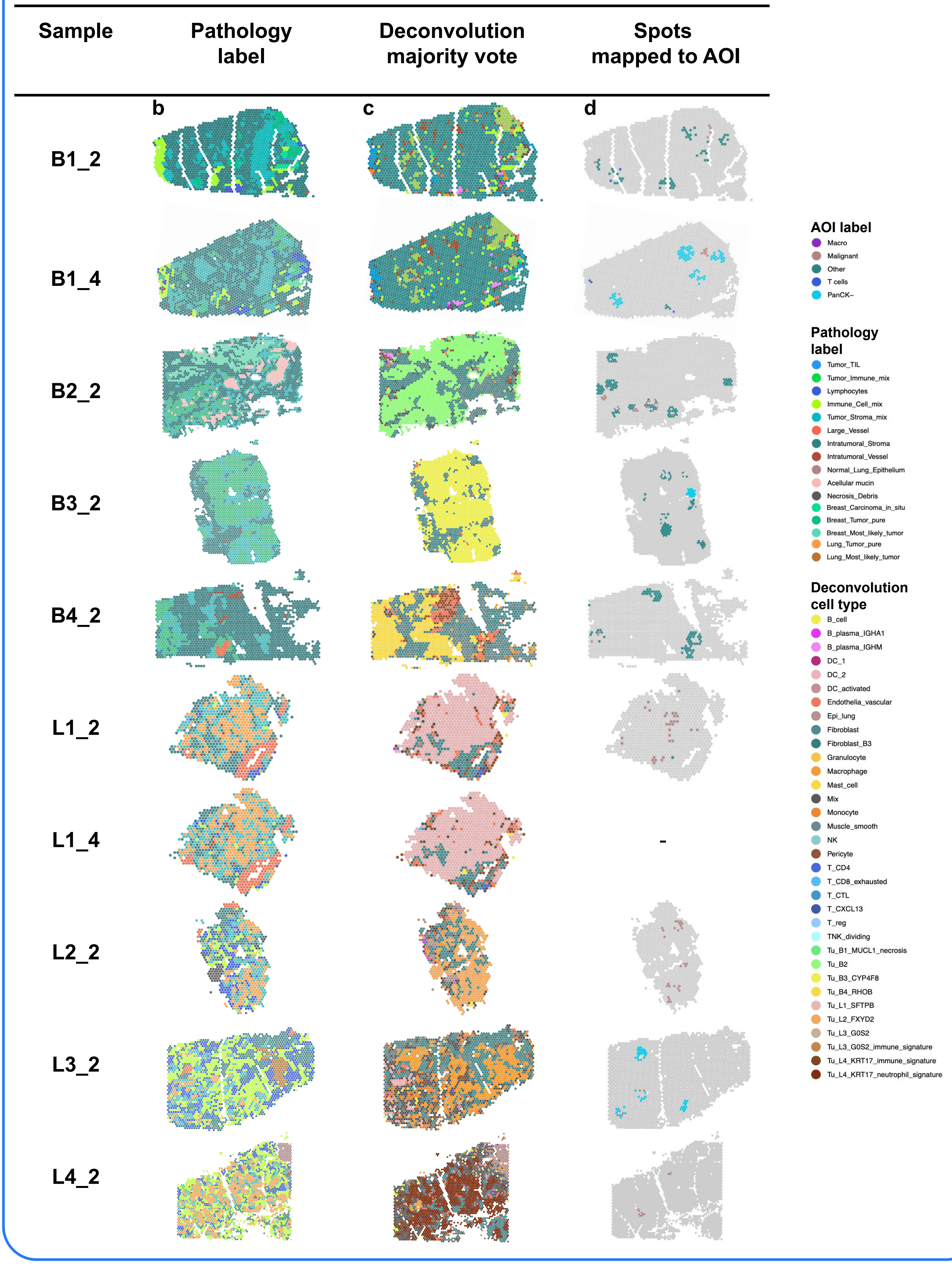

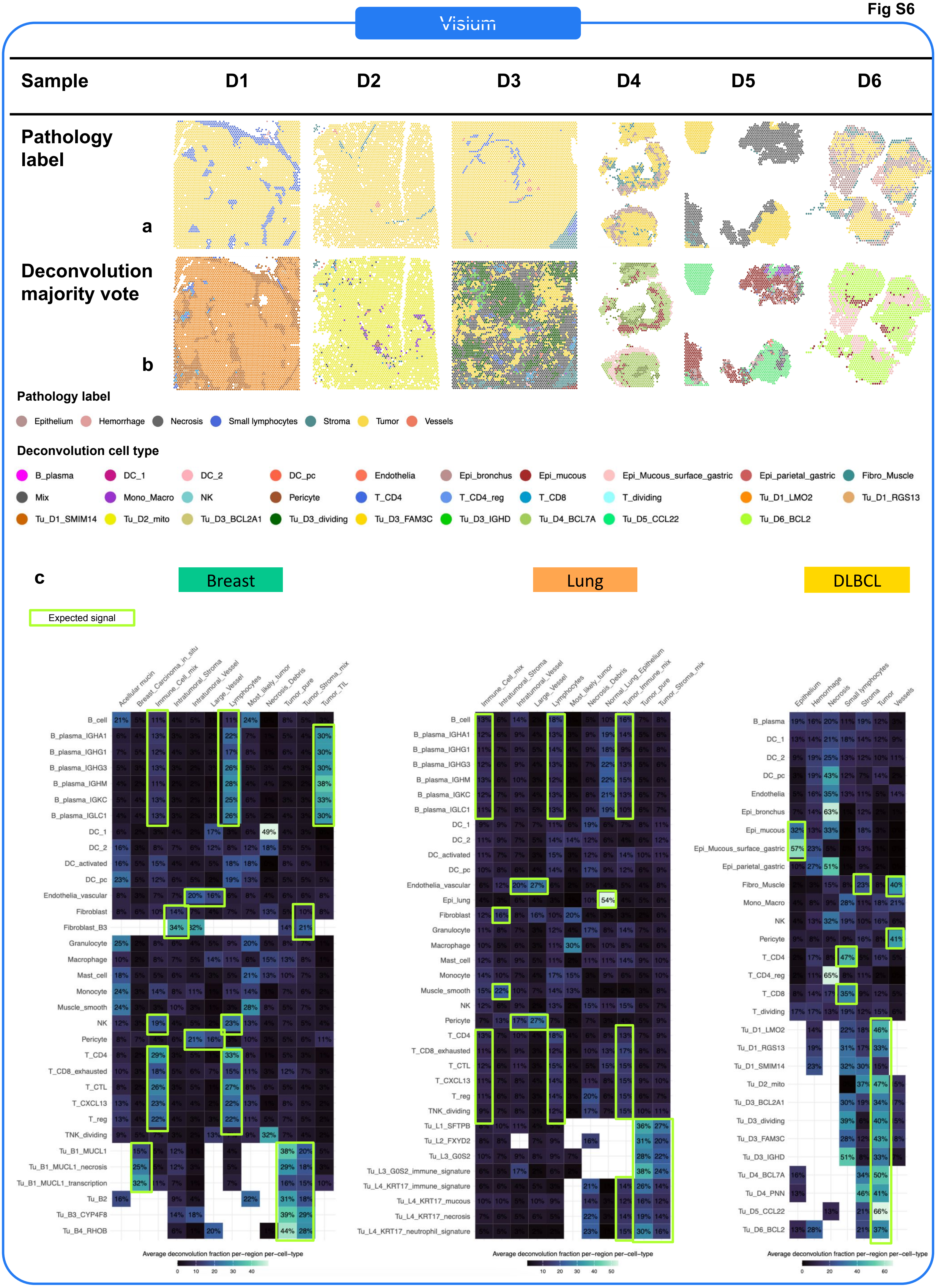

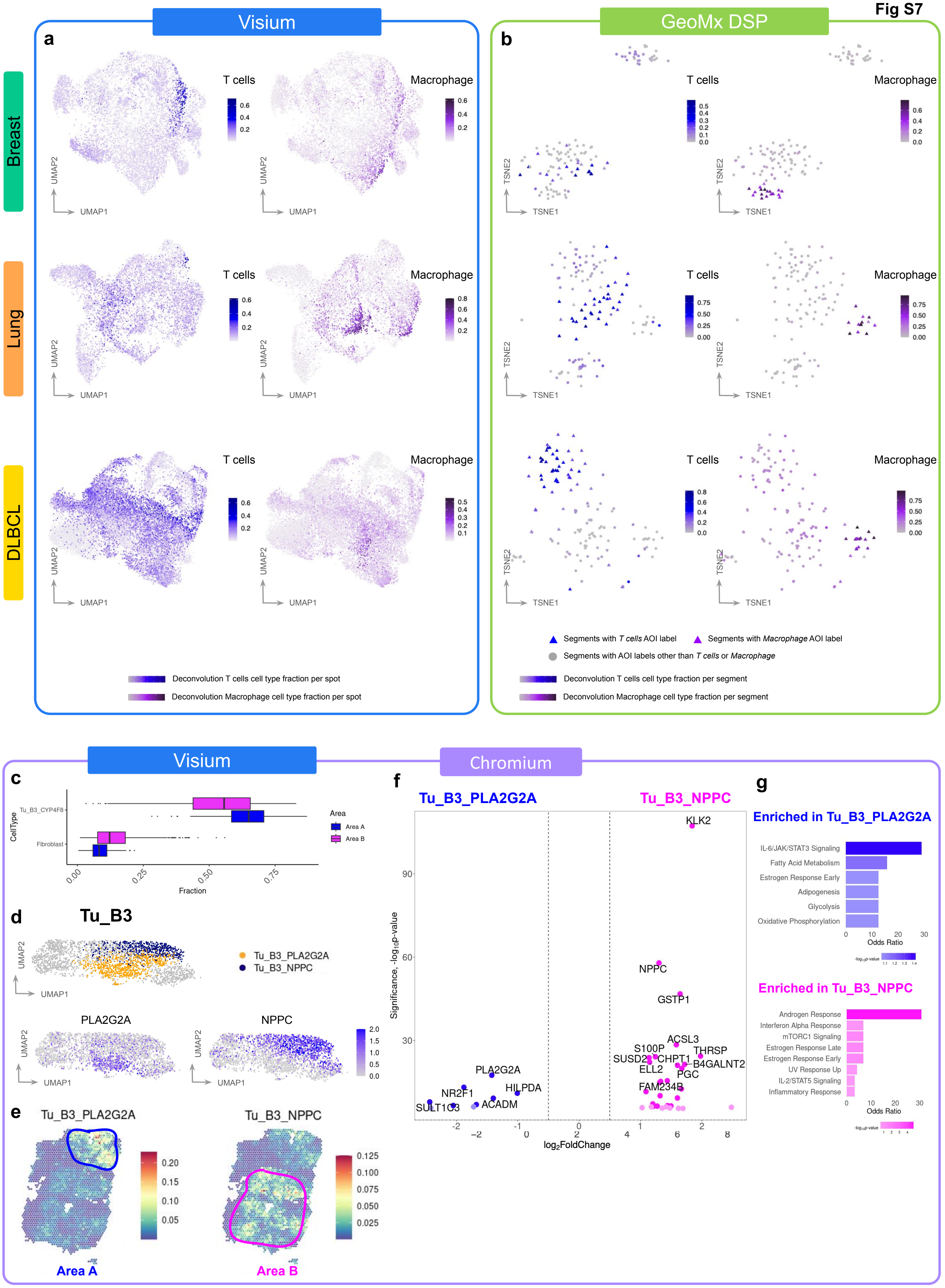

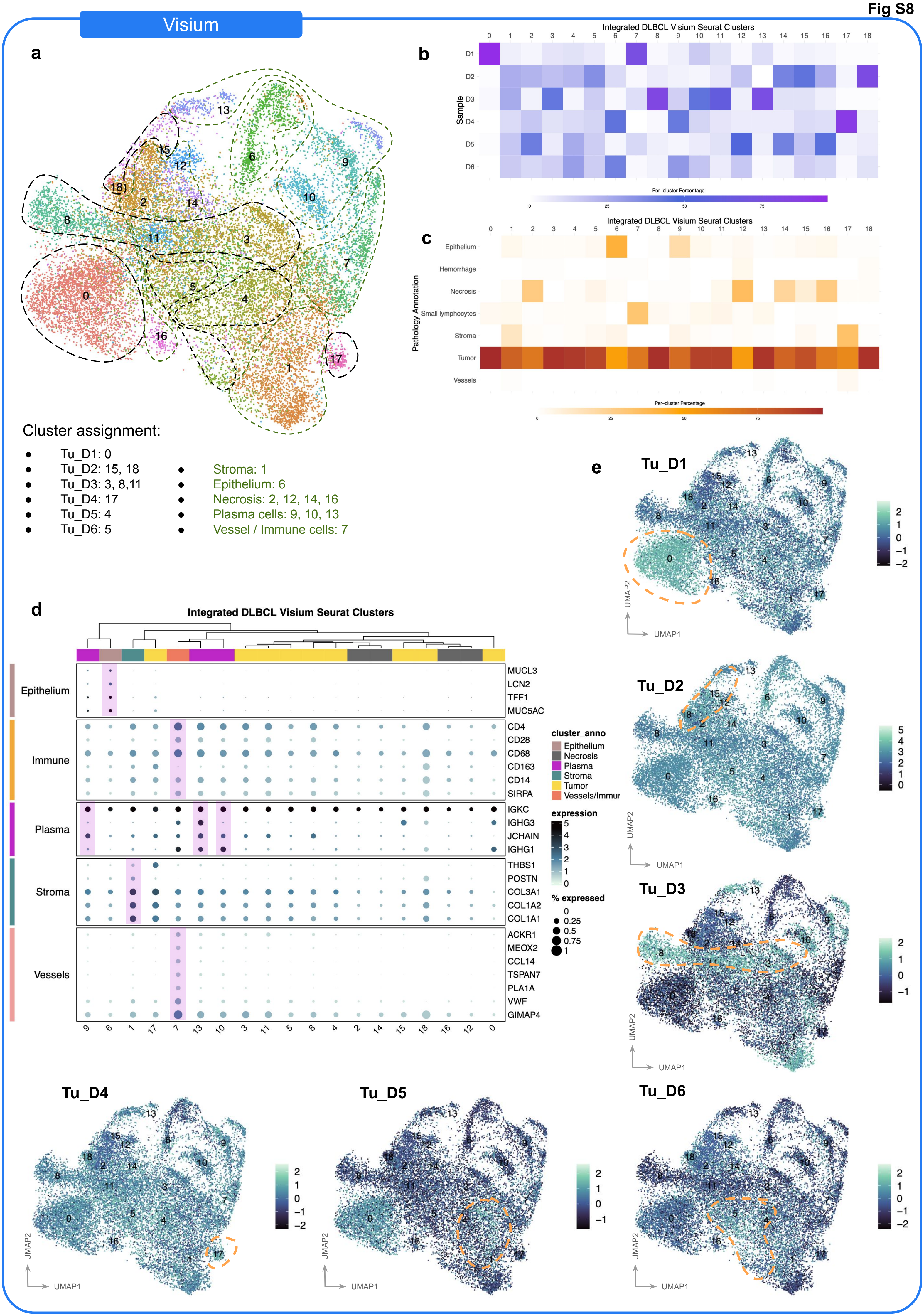

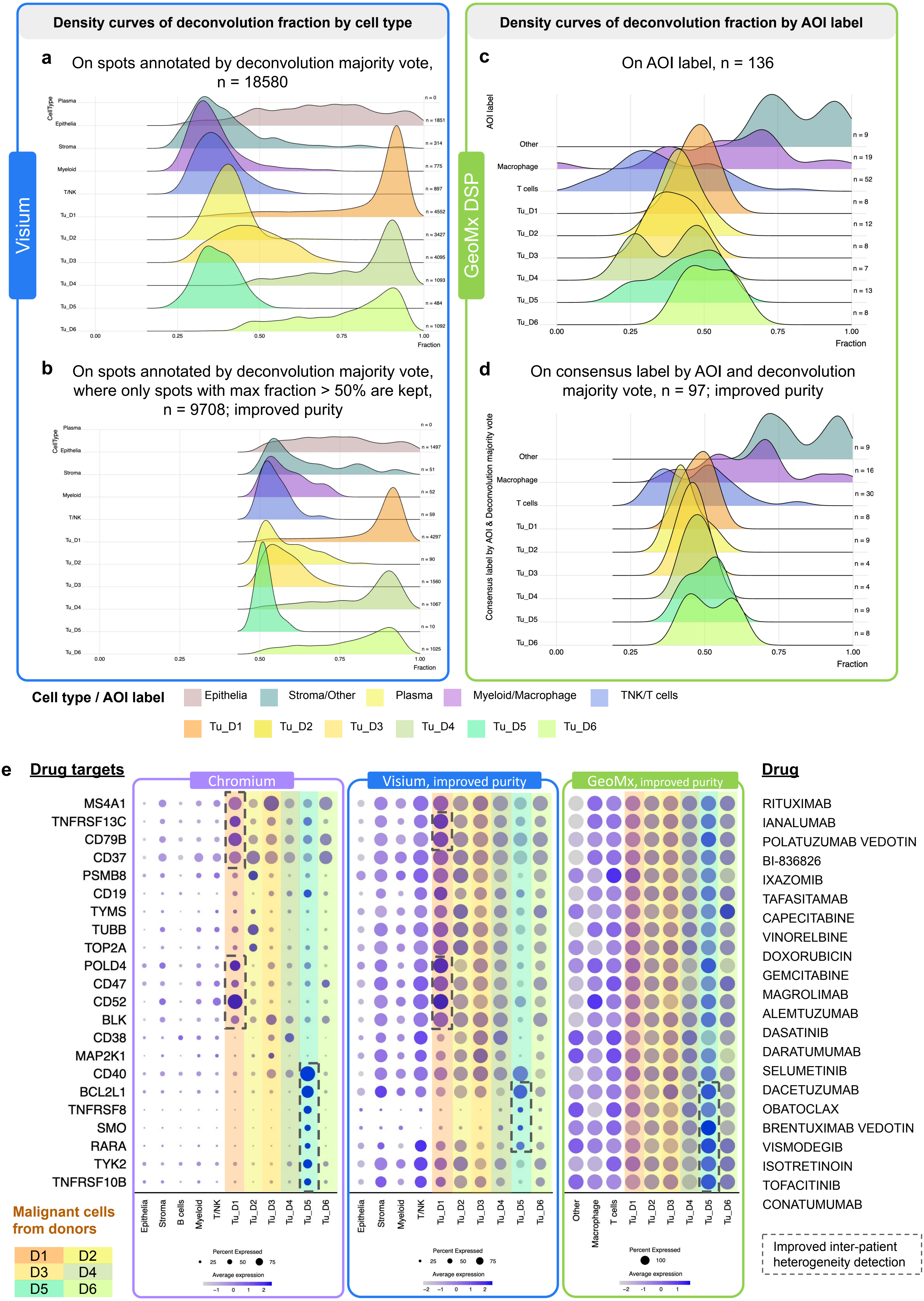
